# GWAS hints at pleiotropic roles for *FLOWERING LOCUS T* in flowering time and yield-related traits in canola

**DOI:** 10.1101/539890

**Authors:** Harsh Raman, Rosy Raman, Yu Qiu, Avilash Singh Yadav, Sridevi Sureshkumar, Lauren Borg, Rohan Maheswaran, David Wheeler, Ollie Owens, Ian Menz, Sureshkumar Balasubramanian

**Author notes:** **Corresponding author:** Dr. Harsh Raman, NSW Department of Primary Industries, Wagga Wagga Agricultural Institute, Wagga Wagga, NSW 2650, Australia.

## Abstract

Transition to flowering at the right time is critical for local adaptation and to maximize seed yield in canola, which is an important oilseed crop. There is extensive variation among canola varieties in flowering time. However, our understanding of underlying genes and their role in canola productivity is still limited. We reveal natural variation in flowering time and response to photoperiod in a diverse GWAS panel (up to 368 accessions) of canola and identify associated SNPs across multiple experiments. Complementary QTL and eQTL mapping studies were also conducted in an Australian doubled haploid (DH) population for flowering time and other grain yield related traits. We show that several associations that were repeatedly detected across experiments map in the vicinity of *FLOWERING LOCUS T* (*FT*) paralogues and its known transcriptional regulators. QTL mapping study in a DH population detected consistent genomic regions close to *FT* paralogs. *FT* sequences vary between accessions and *FT* expression in field and controlled environment grown plants was correlated with flowering time. *FT* paralogs displayed association not only with flowering time, but also with plant emergence, shoot biomass and grain yield. Our findings suggest that *FT* paralogs not only modulate flowering time but also modulate yield-related productivity traits in canola.

**Highlight:** The genetic association, eQTL and expression analyses suggest that *FT* paralogs have multifaceted roles in canola flowering time, plant development and productivity traits.

## Introduction

Natural variation provides a valuable resource to discover genetic and molecular basis of phenotypic diversity in plant development, adaptation and productivity (Alonso-Blanco *et al.*, 2009; Pin and Nilsson, 2012). Canola (rapeseed, *Brassica napus* L., A_n_A_n_C_n_C_n_ genomes, 2n = 4× =38) is an important oil crop, varieties of which displays extensive variation in life history traits such as flowering time. Precise knowledge of flowering time is fundamental for both identifying varieties that are locally adapted and for the development of varieties that are suitable to changing environments, while maximizing grain yield, oil content and quality. Early flowering varieties are preferred for cultivation for shorter season especially under water-limited conditions to escape from excessive drought and heat, whereas winter/semi-winter crops are targeted for longer season under temperate regions to achieve maximum yield.

In *Arabidopsis thaliana*, four major pathways involved in flowering time; photoperiod, vernalisation, autonomous flowering and gibberellic acid response are reported (Koornneef *et al.*, 2004; Weigel, 2012). In addition, flowering is also affected by other external factors such as ambient temperature, insect-pests, pathogens, light quality, and abiotic stress, and some of these integrate with the flowering pathways. Genetic analyses based on classical linkage mapping (quantitative trait loci: QTL) and genome-wide association studies (GWAS) have revealed that flowering time in canola is a multi-genic trait (Ferreira *et al.*, 1995; Long *et al.*, 2007; Nelson *et al.*, 2014; Raman *et al.*, 2016b; Raman *et al.*, 2013; Raman *et al.*, 2016c; Schiessl *et al.*, 2015; Xu *et al.*, 2016; Yi *et al.*, 2018). Candidate genes underlying flowering time variation due to vernalisation have been identified in *B. napus* (Fletcher *et al.*, 2015; Hou *et al.*, 2012; Raman *et al.*, 2016b; Raman *et al.*, 2013; Tadege *et al.*, 2001; Wang *et al.*, 2011; Zou *et al.*, 2012). We have previously shown that *BnFLC.A02* accounts for the majority (∼23%) of variation in flowering time among diverse accessions of canola (Raman *et al.*, 2016b). Nevertheless, little is known about functional role of the photoperiod responsive genes in modulating flowering time especially in spring canola varieties.

*FLOWERING LOCUS T* (*FT*) is a floral integrator and as such generally considered downstream of the photoperiod pathway, integrating inputs from different pathways. In *A. thaliana,* loss-of-function mutations in *FT* gene result in late flowering under long-day conditions (Koornneef *et al.*, 1998; Koornneef *et al.*, 1991). In *B. napus*, six paralogues of *FT* have been identified (Wang *et al.*, 2012; Wang *et al.*, 2009) which contribute to functional divergence affecting flowering time between winter and spring cultivars. Mutation in *BnC6.FTa and BnC6.FTb* paralogs have been shown to alter flowering time in *B. napus* accessions (Guo *et al.*, 2014). Owing to the multiple copies of *FT* in canola, it has been difficult to establish the functionality and precise relationship between various paralogs in plant development and productivity traits, as shown in Arabidopsis, onion and potato (Kinoshita *et al.*; Krieger *et al.*, 2010; Lee *et al.*, 2013; Lifschitz *et al.*, 2006; Navarro *et al.*, 2011; Shalit *et al.*, 2009). In addition, under field conditions, it is difficult to determine the extent of genetic variation in photoperiod response, as plants undergo series of cold temperature-episodes required for vernalisation.

Here we determine the extent of flowering time variation utilizing a panel of diverse 368 genotypes of canola representing different geographic locations of the world. By GWAS, we identify several underlying QTLs controlling phenotypic variation in flowering time and photoperiod response, estimated as difference in days to flower betweenlong- and short day conditions. We show that the response to photoperiod maps to *FT* paralogues, and their potential transcriptional regulators such as *CIB, CO, CRY2, FVE, MSI, EMF2* and *PIF4*. We complement our findings through QTL analysis in a doubled haploid population. Using plants grown under LD and field conditions, we show that expression levels of *FT* paralogs are significantly associated with flowering time variation across diverse canola accessions. The eQTL analysis for *FT* expression levels map not only to *FT* itself (e.g., *BnA7.FT*) but also other loci that are known regulators of *FT* such as *BnFLC.C3b* (*FLC5*), *FPA, SPA1* and *ELF4*. We also demonstrate that plant productivity traits such as plant emergence, shoot biomass accumulation, plant height, and grain yield map in the vicinity of *FT*. Taken together our findings suggest that *FT* has multifaceted role in plants and could be exploited for selection of canola varieties for improved productivity.

## MATERIALS AND METHODS

### Plant material and growth conditions

#### Evaluation of GWAS panel

A diverse panel of 368 accessions of *B. napus* L. was used to evaluate photoperiod response in this study (Supplemental Table S1). A subset of these, 300 accessions were evaluated for flowering time and grain yield (a) in field plots (35°03’36.9”S 147°18’40.2”E, 147 m above sea level) and (b) in single rows (35°02’27.0”S 147°19’12.6”E) at the Wagga Wagga Agricultural Institute (WWAI) research farm located at Wagga Wagga, NSW, Australia (c) at Condobolin, NSW, Australia (33. 0418.98°S, 147.1350.16°E, 220 m above sea level) in 2017canola growing season. For Wagga field trial, 300 accessions were arranged in a randomized complete block design with 60 rows by 10 columns (ranges) in four flood irrigation bays, each bay had 15 rows and 10 ranges (Supplementary Table S2). A buffer row of an Australian canola variety, Sturt TT was seeded after every two ranges to ensure that plots are harvested at the ‘right maturity time’ with a mechanical plot harvester. For Wagga single row trial, 300 accessions were arranged in a randomized block design with 60 rows (each row 10 M long) by 10 columns in two replicates (Supplementary Table S2), each replicate of 30 rows was separated with a buffer row of SturtTT canola variety. The Condobolin trial was arranged in a random complete block design with 100 rows by 6 columns, accommodating all 300 accessions in two replicates (Supplementary Table S2). For field plot experiments, accessions were sown in plots (2 m wide × 10 m long at Wagga Wagga and 2 m wide × 12 m long at Condobolin) at density of 1400 seeds/20 m^2^ plot. Seeds were counted with Kimseed machine and directly sown in plots in the field; each plot consisted of 6 rows spaced 25 cm apart. Plots were sown with a six-row cone-seeder to 10m length. All plots were sown with a granular fertilizer (N : P : K: S, 22 : 1 : 0 : 15) applied at 150 kg ha^−P^. The fertilizer was treated with the fungicide Jubilee (a.i. flutriafol at 250 g Lat 2Farmoz Pty Ltd (Adama), St Leonards, NSW) to protect all genotypes against the blackleg fungus, *Leptosphaeria maculans*. After crop establishment, plots were trimmed back to 8 m after emergence by applying Roundup (a. i. glyphosate) herbicide with a shielded spray boom.

For controlled environmental cabinets (CE), eight plants were grown in plastic trays as described previously (Raman *et al.*, 2016b) under long (LD) and short day (SD) conditions. For LD treatment, seeds were planted in a CE maintained at 20 ± 1°C under white fluorescent lamps (4000 K, Osram) with light intensity of approximately 150µM/m^2^/s, with a 16-h photoperiod. In SD treatment, plants from 368 accessions were grown at the same conditions described above but for 8 h photoperiod.

### Flowering time and other phenotypic measurements

Days to flower were recorded when 50% of plants have opened their first flower from the day of sowing. In SD conditions, flowering time was recorded for up to 200 days. Plants without any flower at the end of the experiments were assigned as value to 200 days; those phenotypes were classified as flower at (LD-A, SD-A, see Fig. 1). The response to photoperiod was calculated as the difference between 50% flowering in plants grown under SD and LD conditions. For field trials, flowering time was recorded three times in a week.

**Fig. 1.**
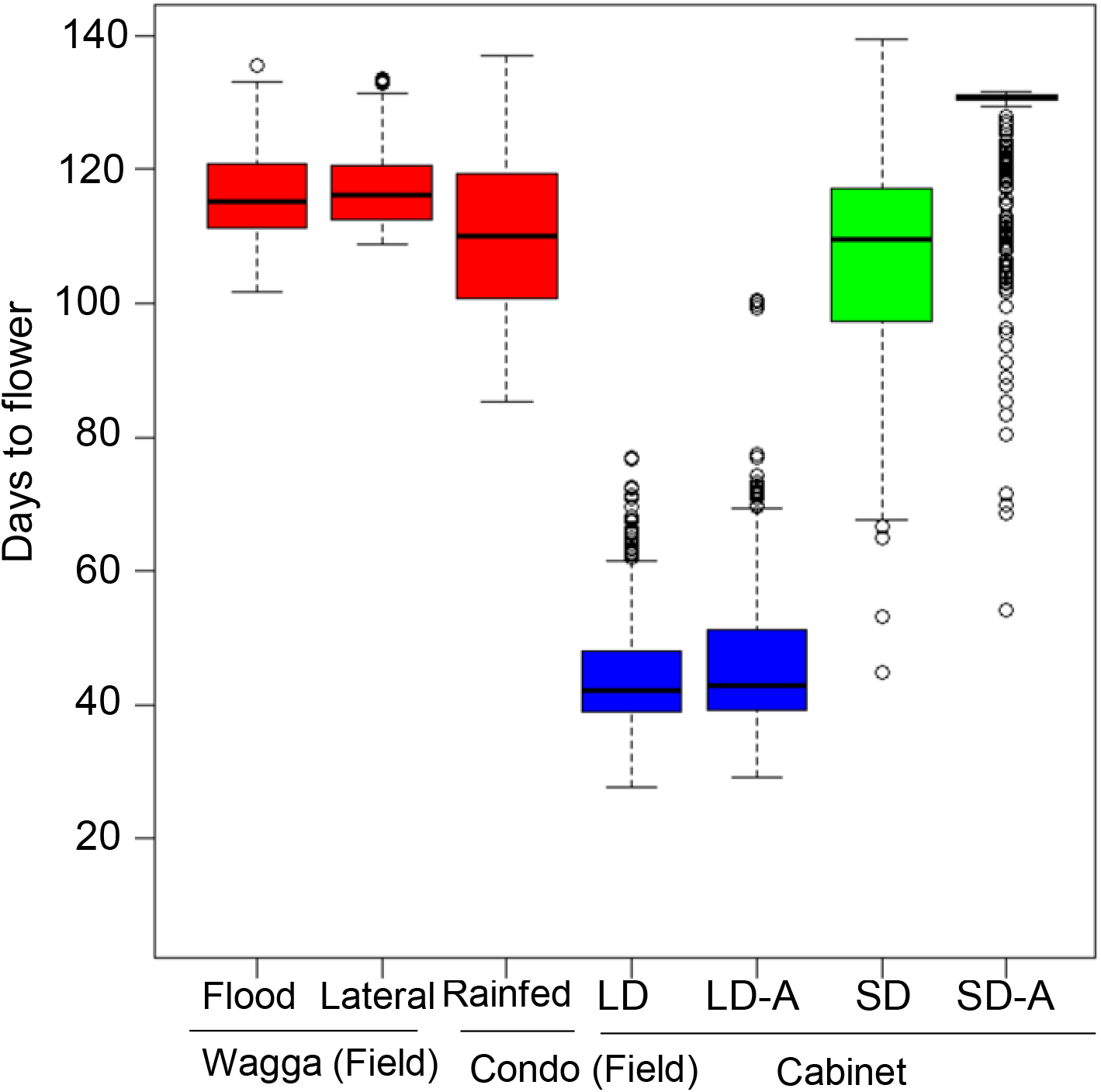
Natural variation for flowering time. Box-plots showing genetic variation for flowering time in a diverse panel of canola accessions grown across five experiments under field, and controlled environment cabinets (cabinets). Days to flowering were scored in 2016 (Field plots at Wagga Wagga (flood irrigated) and Condobolin (Condo); Single rows at Wagga site under lateral move irrigation); Days to flowering were scored in long day condition (LD, 16 h) and short day condition SD) plants under cabinets. Black dots indicate genotypes that did not flower till the end of experiment and were marked as ‘assigned’ (LD-A and SD-A). A total of 368 accessions were evaluated for flowering time under LD and SD conditions, while 300 accessions were evaluated under field conditions. Details are given in Supplementary Table 1.

Normalised Difference Vegetative Index (NDVI) was measured as a proxy of fractional ground cover for early vigour (Cabrera-Bosquet *et al.*, 2011; Cowley *et al.*, 2014) using a GreenSeeker® hand-held optical sensor unit (model 505, NTech Industries Inc., Ukiah, CA, USA). The NDVI readings were taken at 7-10 days interval after 5 weeks of sowing before the onset of flowering. Multiple readings were taken in each plot and then averaged across each plot for genetic analysis. Plots were harvested by direct heading with a Kingroy plot harvester (Kingaroy Engineering Works, Queensland, Australia) in the 4th wk of November (Condobolin, NSW) and 2-3 wk of December (Wagga, Australia). Grain samples were cleaned with Kimseed (Kimseed Australia, Western Australia) and plot yield was expressed into t/ha.

### Field evaluation of SAgS DH population

A population of 144 DH lines derived from a BC_1_F_1_ plant between Skipton/Ag-Spectrum//Skipton; SAgS DH population (Raman et al 2016) was grown in 2015 (35°01’32.3”S 147°19’25.4”E) and 2016 (35°01’42.8”S, 147°20’23.3”E) in the field at the WWAI, NSW, Australia. Both trials were randomized in a complete block design with three replicates in a single block. A total of 1,400 seeds per genotype were directly sown in plots in the field as described above. The number of traits were measured, including plant emergence, first flowering, plant biomass, plant height, and grain yield. Plant (shoot) biomass was measured by cutting 10 random plants from the one meter central row of each plot and then brought back to the canola shed. Each sample was weighed on a digital scale and fresh weights were expressed in g/plant. Plant height (cm) was measured at the physiological maturity stage by measuring 5 plants selected randomly in the middle row of each plot. Plots were harvested with a Kingaroy plot harvester in the 2-3 wk of December (Wagga, Australia).

### Genome-wide genotyping

Leaf material was collected from 368 diverse DH canola accessions, grown under LD conditions, and then immediately snap-frozen in liquid nitrogen. Genomic DNA was isolated following method described previously (Raman et al 2016) and sent to Trait Genetics, Germany (www.traitgenetics.com/) for genotyping with Illumina infinium 15k *Brassica* chip representing 60K Infinium SNP array (Clarke *et al.*, 2016). Markers which have the overall call rate over 90% were used for trait-marker association analysis. To prevent the potential loss of GWA, missing data was imputed (Rutkoski *et al.*, 2013).

### Population structure and GWA analyses

The SNP markers with allele frequency <0.05 and call rate <90% (Atwell *et al.*, 2010) from the 13,714 genome-wide SNPs, were discarded before GWA analysis. Of them, 11,804 SNP markers could be anchored to the A_n_ and C_n_ subgenomes of reference sequenced genome of *B. napus cv. ‘*Darmor-*bzh’,* hereafter Darmor and used for cluster, and GWA analyses in a diversity panel of 368 accessions (S1 Table). Cluster analysis was performed with Neighbor-Joining method (Saitou and Nei, 1987) in MEGA version 6. In order to reduce spurious associations between markers and variation in flowering time, population structure and the relative kinship coefficients of individual genotypes were estimated as described previously (Raman *et al.*, 2016b). Flowering time-SNP marker association analysis was performed using the EMMAx/P3D method (Kang, 2008; Zhang *et al.*, 2010) implemented in GAPIT (Lipka *et al.*, 2012) in R package (http://cran.r-project-org). Significance of GWA between markers and flowering time was tested at LOD score of 3. The *P* (–log_10_*P*) values for each SNP were exported to generate a Manhattan plot in R (Team, 2014). The proximity of candidate genes to identified associations based on the physical positions of SNPs/candidate genes was inferred based on functional annotation of the *A. thaliana* genome and implemented in the reference sequenced genome of *‘*Darmor’ (Chalhoub *et al.*, 2014). Using Bonferroni correction, associations with LOD score = 5.41) were also considered as significant on a *p*<0.05 level. The associations detected through GWAS, were compared against the QTL marker intervals associated with flowering time under field conditions in a SAgS DH mapping population evaluated in 2013 and 2014 (Raman *et al.*, 2016c) and in 2015 and 2016 (this study).

### Statistical and QTL analysis

Flowering and other phenotypic data collected from different experiments were analysed using linear mixed models in R as described previously (Raman *et al.*, 2018). Essentially we defined the individual experimental Plot as factor, with 432 levels for each of the 2015 and 2016 trials. The factors Row and Range corresponded to the rows and ranges of the trials, with levels equal to the number of rows and ranges in each trial. The combination of levels of Row and Range completely index the levels of Plot such that Plot = Row:Range. The factor Rep has 3 levels corresponding to the replicate blocks in each trial. The plot structure for the field experiment consists of plots nested within blocks and is given by, Rep/Plot which can be expanded to give, Rep + Rep:Plot The term Rep:Plot indexes the observational units for all traits and so is equivalent to the residual term for these traits. The treatments for the field phase of the experiment are the lines allocated to plots and so we define the treatment factor, Genotype, with 144 levels corresponding to lines grown in each trial. Due to marker data being included in the model, we need to define an additional two factors; Gkeep (corresponding to lines with both phenotypic and marker data) and Gdrop. factor Gdrop has 16 levels corresponding to lines with phenotypic data but not marker data. Therefore treatment stucture is given by, Gkeep + Gdrop. Finally, marker data is incorporated into the analysis and individual markers are scanned following the approach of Nelson et al. (2014) to establish a final multi-QTL model. We also used phenotypic data from 2013 and 2014 experiments that was published previously (Raman *et al.*, 2016c), in order to test multifaceted role of *FT* in flowering time and other productivity traits across environments. Genetic map based of 7,716 DArTseq markers representing 499 unique loci (Raman *et al.*, 2016c) was used to determine trait-marker associations. The predicted means for first flowering, and response to photoperiod for each genotype were used to detect genome wide trait-marker associations.

### *FT* expression and eQTL analyses

For the of *FT* expression analysis from extreme phenotypes, 24 accessions were selected from 368 GWAS accessions based of their flowering time and photoperiodic response. These accessions were raised in LD conditions under CE cabinets as described above using an experimental design with four replications and scored for flowering time (Supplemental Table S2). Five independent leaf samples from field/CE grown plants (at floral budding stage) per genotype; 24 GWAS and 144 DH lines of SAgS DH mapping population, were pooled and flash-frozen in liquid nitrogen (in field/CE).RNA was isolated using TRIZol (Invitrogen) and cDNA was synthesized using First Strand Synthesis Kit (Roche). Samples were controlled for their quality using two different approaches as outlined previously (Raman *et al.*, 2016b). The gene specific primers for each of six *FT* paralogs (Guo *et al.*, 2014) were used for the expression analysis (Supplemental Table S3). Since all *FT* paralogs showed a high correlation among themselves, we used *BnC6.FT* gene expression data for eQTL analysis using SVS package (Golden Helix, Bozeman, USA).

### Structural variation in canola *FT* paralogs

We generated the whole-genome resequence data for the 21 canola accessions (Raman et al, unpublished) representing our GWAS panel including both parental lines; Skipton and Ag-Spectrum, of the SAgS mapping population used in this study (Supplementary Table 2). Data was generated using the Illumina HiSeq 2000 sequencing platform using paired-end reads (150 bp). Reads were mapped on to reference genome assembly (version 4.1) of cv. ‘Darmor’ using BWA (version 0.7.8). SNP and indel calling based on the short read alignment data was performed using the GATK haplotype caller (version 3.5). Variation across the *FT* paralogs was extracted using the gene model information or by manually identifying gene regions based on BLAT homology (Supplemental Table S4). The physical positions of different *FT* paralogs (NCBI GenBank accessions; genomic sequences: FJ848913 to FJ848918; promoter sequences: JX193765, JX193766, JX193767, JX193768) were confirmed with those of the sequenced *FT* genes on the ‘Darmor’ assembly as well as with published literature (Schiessl *et al.*, 2014; Wang *et al.*, 2012; Wang *et al.*, 2009). For each accession, the *FT* nucleotide sequences were aligned using MUSCLE as implemented (Edgar, 2004) in the software package, Geneious (https://www.geneious.com) and estimated for structural variation, number of polymorphic sites in exons, intron, and promoter regions using ANNOVAR (Wang *et al.*, 2010). The diversity indices were calculated using the MEGA version 6 (Tamura *et al.*, 2013). The Tajima (1989) and Fay and Wu (2000) tests were conducted to examine whether the frequency spectrum of polymorphic nucleotide mutations conformed to neutral expectations. The effect of InDel mutations on functional domains was investigated using information from the NCBI conserved domain database.

## RESULTS

### Natural variation in flowering time across diverse environments

We determined the natural variation in flowering time of diverse accessions in two different environmental conditions across five separate experiments. Across all phenotypic conditions, we found extensive variation in flowering time, which ranged from as early as 27.6 days up to more than 139.4 days (Fig. 1, Supplemental Table S5-6). Diverse accessions grown under LD conditions (16 h light at 20°C) in controlled conditions typically flowered earlier (27.6 to 77 days) compared with SD conditions (44.9 to 139.4 days under 8 h light at 20°C in growth cabinet) and accessions grown in field conditions (85.2 to 137.1 days). Accessions grown under rainfed conditions (Condobolin site) flower earlier compared to irrigated sites (Wagga) Supplemental Table S6. Most of this variation was genetically controlled as the broad sense heritability (*h^2^*, also called as reliability) ranged from 45% to 97% across different environments (Supplemental Table S7). We observed positive genetic correlations (*r* = 0.88 to 0.96) for flowering time between the different field trials, suggesting that majority of the genetic variation and underlying mechanisms are shared across field environments (Fig. 2).

**Fig. 2.**
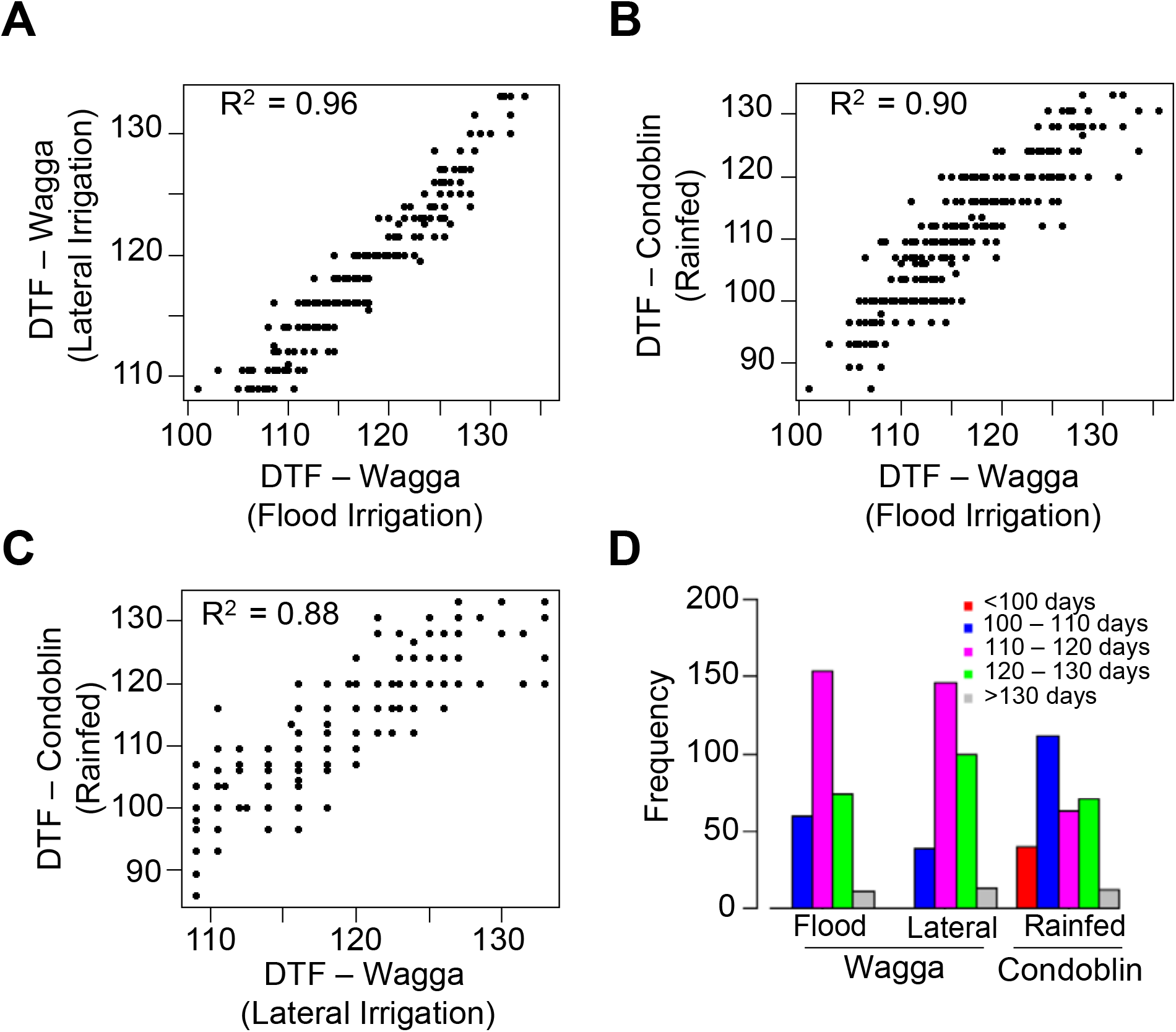
Pearson correlation for flowering time among 300 accessions of canola evaluated in field plots across different environments. Flowering time (days to flower, DTF) was assessed thrice in a week. A) Flowering time correlation between field trials that were irrigated with lateral move or via flooding. B) Flowering time correlation between field trials at flood irrigated plots at Wagga with rainfed plots at Condobolin. C) Flowering time correlation between laterally irrigated plots at Wagga and rainfed plots at Condobolin and D) Frequency distribution of canola accessions based on the days to first flower under the varied conditions.

### Flowering time variation in canola is largely due to photoperiodic response

Under controlled environmental conditions in growth cabinets, LD photoperiod substantially promoted flowering (27.6 to 77 days) (S1 Table, Fig. 1), while only 23.8% of accessions (n = 86) flowered under short days, suggesting that extended photoperiod is required for flowering. Analysis of photoperiodic response in accessions enabled us to identify specific accessions of interest, with robust photoperiod sensitive or insensitive behavior (Fig. 1, Supplemental Table S5). Only a small proportion (6.8%, n = 25) of accessions did not flower within 100 days under LD conditions None of the winter type accessions (e.g., 03-P74, Beluga, Ding10, FAN28, FAN168, Gundula, Haya, HZAU-1, Maxol, Rangi, Norin-20, Tower, Zhongshuang-4, Zhongyou 8) either flowered under LD or in SD condition, reconfirming that vernalisation is essential for flowering in those accessions. This is consistent with these genotypes being winter/semi-winter type requiring vernalisation to flower (Raman *et al.*, 2016b).

To assess whether there is any differential photoperiodic response, we compared the effects of photoperiod on flowering time of the accessions grown under controlled environment (CE cabinets). Four accessions, 9X360-310 (BC15278), Georgie (BC15289), CB-Tanami (BC52411) and Hylite200TT (BC52662) had variable response compared to others, suggesting genotype x environment interactions (S1b Table, Supplemental Figure S1).

### Relationship between flowering time and other traits

To determine whether there is any relationship between flowering time and yield-related traits in canola, we calculated Pearson correlation coefficients (Fig. 3). There were low genetic correlations for flowering time between the different field and controlled environmental conditions, suggesting that phenotyping environment play an important role in trait expression. Flowering showed a negative correlation with grain yield across sites (WW-Wagga Wagga and Con: Condoblin) under long day photoperiodic conditions (field and controlled environments). Early vigour (NDVI.WW) showed positive correlations with flowering time (0.2 to 0.7) under LD and field conditions (Wagga and Condobolin), and with grain yield (0.1 to 0.4) depending upon growing environment.

**Fig. 3.**
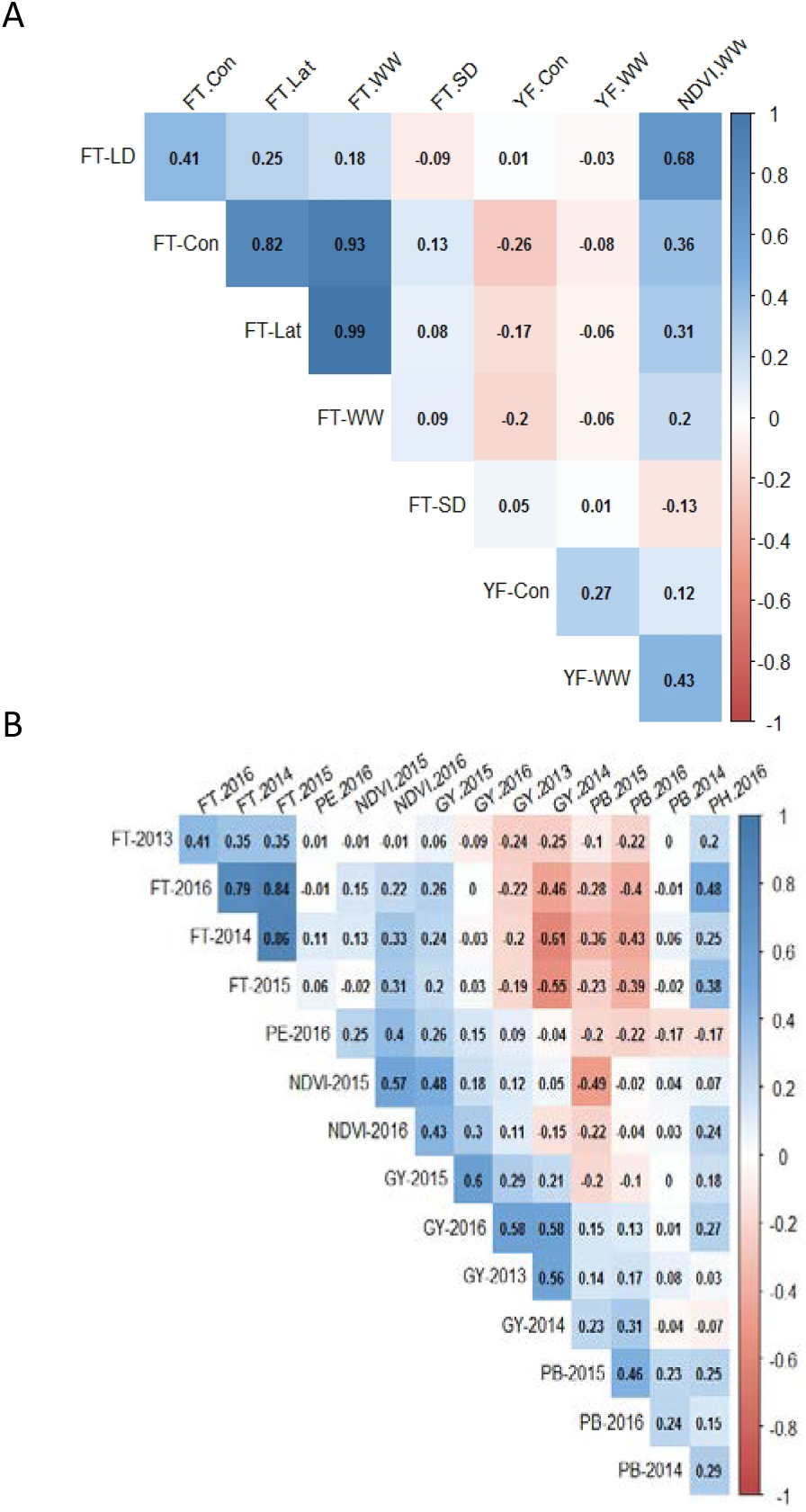
Pearson correlation between flowering time (FT) and yield related traits in a GWAS panel (A) and DH population derived from Skipton/Ag-Spectrum//Skipton (B). FT-LD: flowering time (days to flower under LD conditions; FT-SD: flowering time (days to flower under SD conditions; FT-Con: flowering time at Condobolin; FT-Lat: flowering time at Wagga (lateral move); FT-WW: flowering time at Wagga (rainfed); YF-Con: Grain yield at Condobolin; YF-WW: Grain yield at Wagga (rainfed), NDVI.WW: Normalised Difference Vegetative Index at Wagga; PE: plant emergence; GY: grain yield; PB: plant biomass (g/plant) and PH: plant height (cm).

### Population structure in a GWAS panel

SNP marker distribution across genome is shown in Supplemental Figure S2; mean marker density 621.3 per chromosome provided coverage of ∼84.7 Kb/marker. Cluster analysis revealed that at least three main clades among accessions representing European winter, Australian semi-spring/Canadian spring, and semi-winter of Indian/Chinese origin (Fig. 4, Supplemental Fig. 3). The first three principal components (PC1 = 38.1%, PC2 = 11.9%, and PC3 = 5.67%) accounted for 55.7% of the genetic variation and largely resembled the cluster analysis with similar grouping of accessions (Supplemental Fig. S4). To estimate the extent of genome-wide LD, we calculated the squared allele frequency correlations (average *r*^2^) for all pairs of the anchored SNPs using an LD window of 500 as 0.02 (Supplementary Fig. S5). The VanRaden kinship coefficient among accessions ranged from 0.03 to 0.99 suggesting a wide-range of familial relatedness between pairs of accessions (Supplementary Table S8), as observed in our previous study (Raman *et al.*, 2016b).

**Fig 4:**
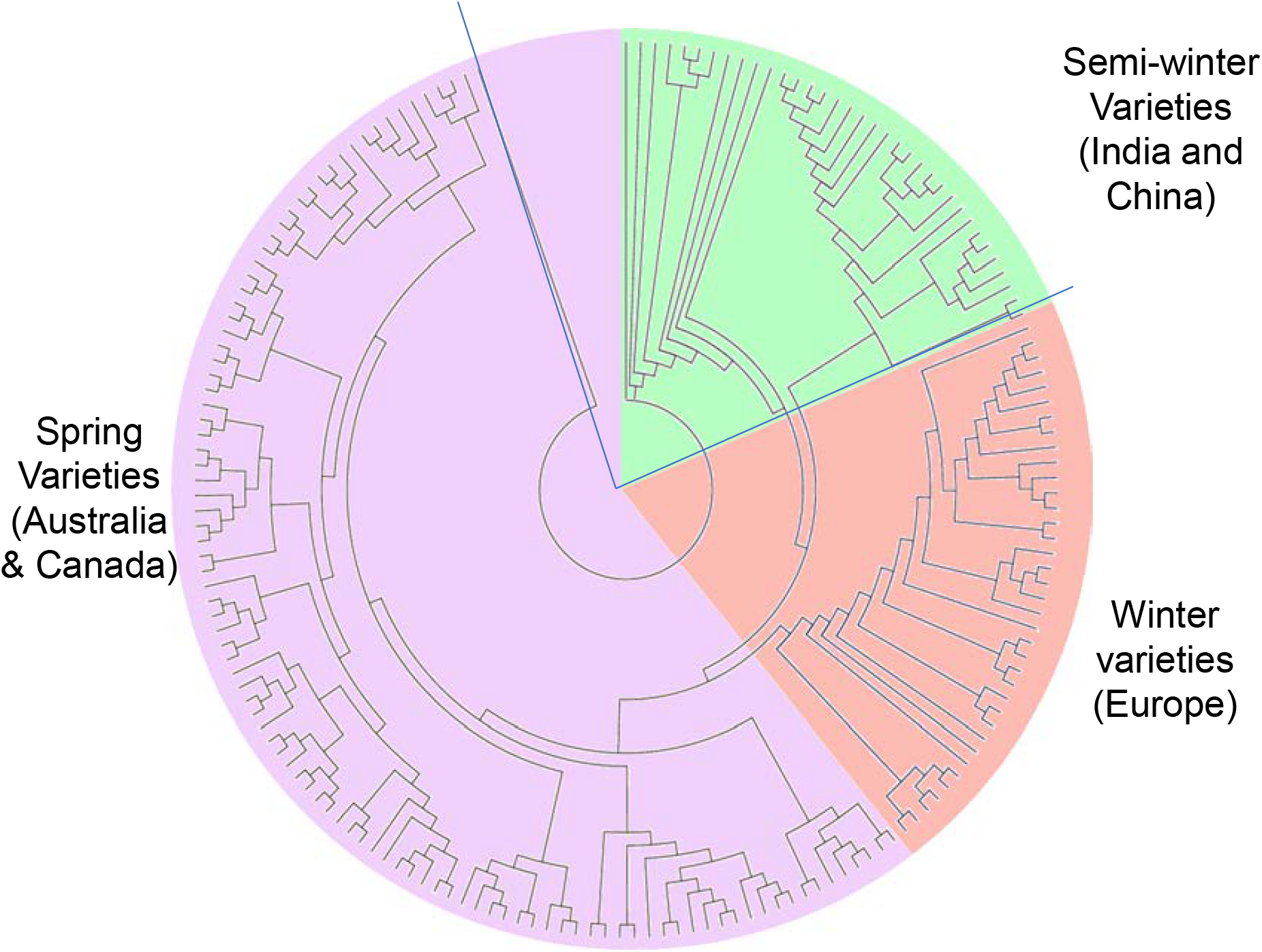
Molecular diversity in a GWAS panel of 368 *Brassica napus* accessions. (A). Three dominant clusters shown in different shades; violet, red and light green colours represent predominantly Australian, European, and Indo-Chinese origins cultivars, respectively. Details are given in supplementary Table x. Tree was drawn with MEGA 6.

### Genetic architecture of flowering time and photoperiod response

Accounting both population structure and kinship information, we identified 142 significant associations (at the genome-wide significance thresholds of LOD score of ≥3) for flowering time under field (three experiments), LD and SD conditions distributed throughout the genome, except on chromosome A01 (Supplemental Table S9). Majority of the associated SNPs (70%) were identified on “A_n_” subgenome (Supplemental Table S10), suggestive of an uneven distribution on the physical locations of ‘Darmor’ assembly. Most of the associated SNPs (33.1%) were on chromosome A02 (47 SNPs) followed by 9.15% on (13 SNPs) explaining majority of allelic variation for flowering time in canola. We identified 22 unique SNP markers that accounted for associations that were detected repeatedly across multiple environments (at least 2 environments, Supplemental Table S9). Of the 142 significant associations, six SNPs crossed the Bonferroni threshold for flowering time in LD conditions, all of which are located on chromosome A02 (Table 1). Two of these SNPs (Bn-A02-p9371948 and Bn-A02-p9371633) associated with flowering time under LD conditions were mapped near the *FT* locus (∼0.64 Mb, *BnA02.FT, BnaA02g12130D*) (Fig. 5A-C). Under different phenotypic conditions, we detected different associations; several of these SNP associations were mapped near the vicinity of genes known to play a regulatory role in *FT* expression in *A. thaliana* such as *FLC4, UPSTREAM OF FLC, CO, MSI1, LD, MAF4* on A02; *BnFLC3a, CO* and *EMF2* on A03; *NY-YB8* on A04; *GI* on A08; *EMF2* and *CRY2* on A10, and *CIB1* on C08 (Supplemental Table S11). We identified 28 SNPs that showed significant association above a LOD of 3 with response to photoperiod identified under controlled environment cabinet conditions on chromosomes A01, A02, A07, A09, A10, C01, C03, C06, C08 and C09 (Supplementary Table 11, Fig. 5C), suggesting that these associations truly reflect genetic determinants of photoperiod response.

**Fig. 5.**
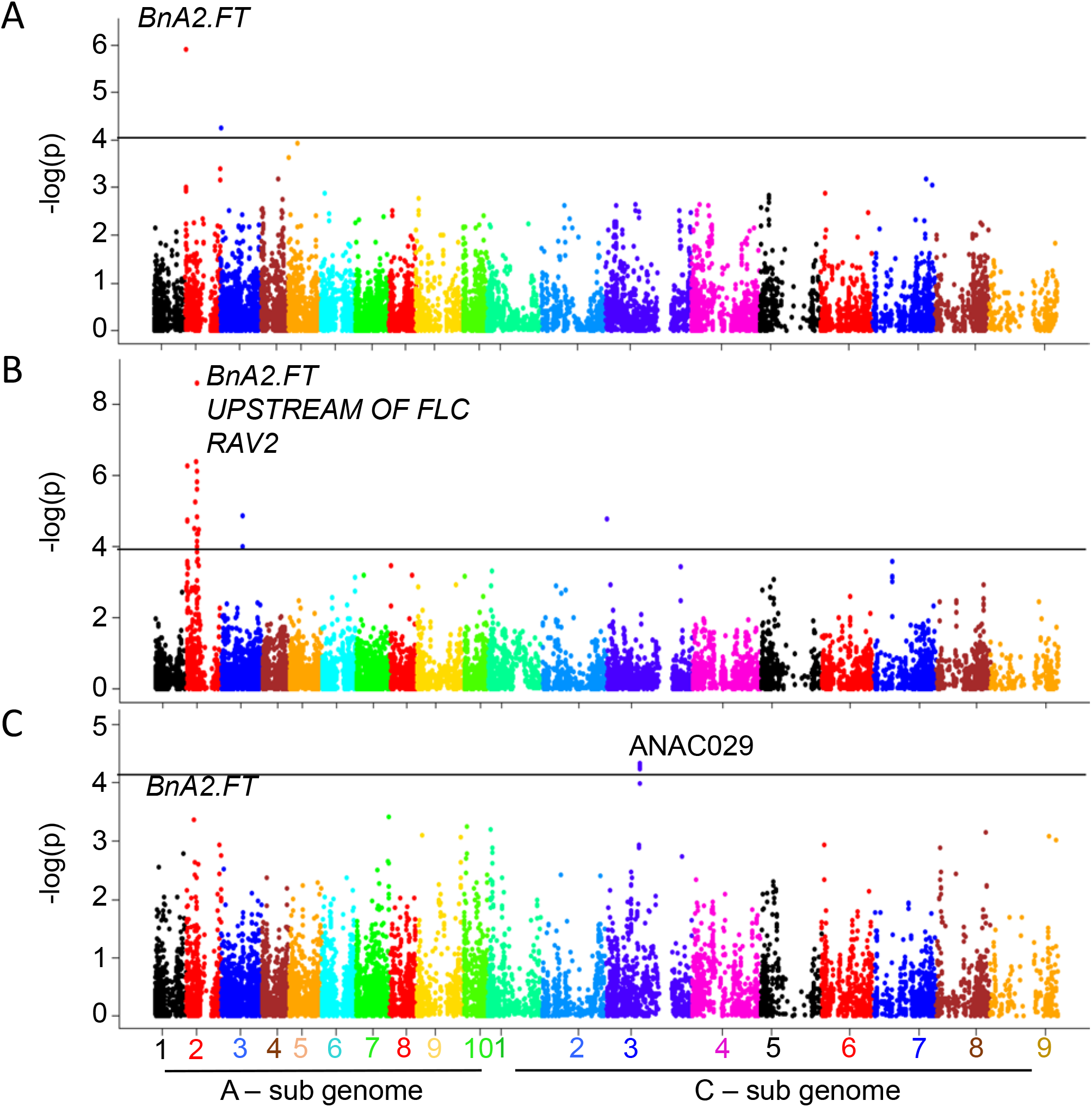
Manhattan plots for the detected associations for flowering time. Plots show genome-wide *P* values for associations between SNP markers and flowering time: (A) Field condition at Condobolin, Australia, (B) long-day conditions in controlled environnent cabinet and (C) response to photoperiod. Different colours represent different chromosomes of *B. napus* (A1-A10, C1-C9). Significant associations - log10(*p*) value of ≤ 4 are shown with a solid horizontal line (in black colour).

To identify potential candidates involved in the photoperiod response, we compared the physical positions of 28 significant SNP associations for photoperiod with the physical positions of flowering time genes (Supplemental Table S11). Of them, seven SNP markers map in the vicinity (0.2 Mb) of *SPA3* (A01), *PRR5* (A02), *MAF4* (A02), *ASH1* (A07), *POWERDRESS* (A10) and *ELF6* (C09), genes underlying photoperiod response in canola accessions; of which ANAC029, *EFF6, ABF2, FVE*, and *PAF1* were detected in CE experiments and *ANAC029*, and *ASH1*, were detected (within 200 kb) under field experiments (S11 Table). Consistent with our previous study (Raman et al 2016a), our results reinforces that while the major players of flowering time appear to be conserved between Arabidopsis and canola, the specific roles of the paralogs might be different depending on the environmental conditions.

### QTL analysis in biparental population identifies loci for flowering time and productivity traits near *FT* paralogs

To ensure capturing the relevance of entire genetic architecture of flowering time variation, we considered the SAgS DH mapping population derived from a BC_1_F_1_ cross between Australian spring type cultivars; Skipton (less responsive to vernalisation) and Ag-Spectrum (more responsive to vernalisation), which was previously utilised for genetic analyses for range of traits of interest (Luckett *et al.*, 2011; Raman *et al.*, 2016a; Raman *et al.*, 2013; Raman *et al.*, 2016c; Raman *et al.*, 2012; Tollenaere *et al.*, 2012). There were moderate to high genetic correlations for flowering time, early vigour, plant biomass and grain yield across environments (phenotyping years) in the SAgS DH population (Fig. 6). Flowering time showed generally negative correlations with grain yield and plant biomass, whereas it showed positive correlation with early vigour and plant height. We identified several QTL associated with flowering time, plant emergence, shoot biomass, plant height, and grain yield across phenotypic environments in the SAgS population (Supplemental Table S12b).

**Fig. 6.**
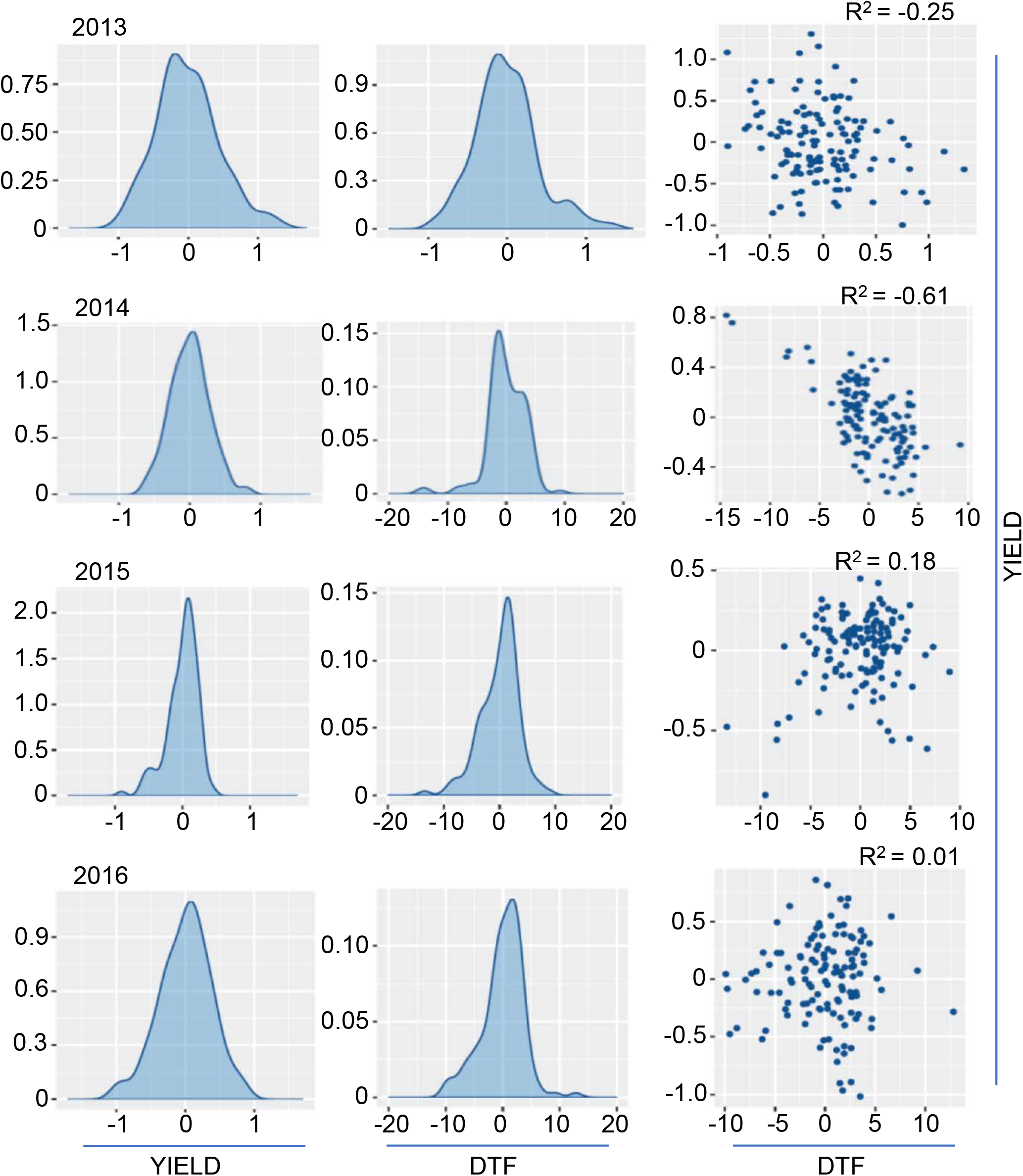
Distribution of flowering time variation in the biparental mapping population. Pair-plots showing genetic correlation of EBLUPS (empirical best linear unbiased estimators) from the univariate analysis of flowering time and grain yield among 144 doubled haploid lines of *B. napus* population derived from Skipton/Ag-Spectrum//Skipton. DH lines were grown across 4 phenotypic environments (2013-2016) in field plots, 2013 at Euberta, NSW, Australia; 2014 at Wagga Wagga, NSW, Australia (Raman et al 2016), 2015 and 2016 at the Wagga Wagga (this study).

Since we detected moderate to high genetic correlations in this population between multiple traits including flowering time (Supplemental Table S13), we considered whether the QTLs underlying these multiple phenotypes co-localise onto the physical map of *B. napus*. Genetic and physical localisation of markers on ‘Darmor’ reference genome (Chalhoub *et al.*, 2014) revealed that three significant QTLs are associated with multiple traits are co-located (Fig. 7). A multi-trait QTL flanked with 3110489 and 3075574 markers for plant emergence, shoot biomass, flowering time, and grain yield mapped on chromosomes A07 was located within 0.65Mb of the *FLOWERING LOCUS T* (*FT*, NCBI accession FJ848914.1); *BnA02.FT* paralog in *B. napus* (Wang *et al.*, 2009). Consistent with GWAS analysis, we detected QTLs near the *FT* in the biparental population (Fig. 7). Mapping of pleiotropic trait QTL in the vicinity of *FT* (A07) suggest that *FT* may have multifaceted role in plant development and productivity traits.

**Fig. 7.**
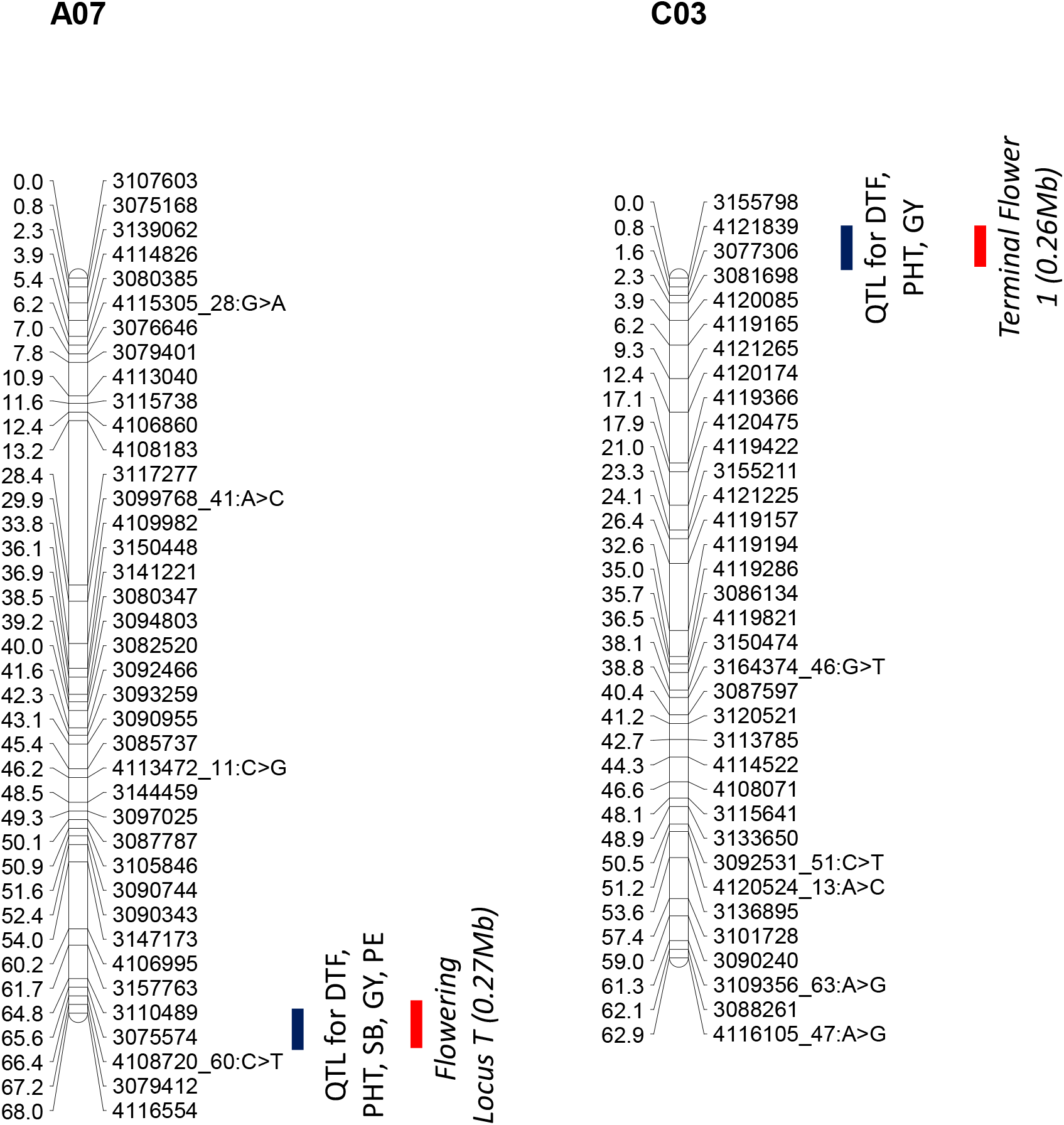
Graphical representation showing localisation of multi-trait QTL associated with plant emergence (PE); above ground shoot biomass (SB); flowering time (days to flower, DTF); plant height (PHT) and grain yield (GY) in a doubled haploid population from Skipton/Ag-Spectrum//Skipton. DArTseq markers and their genetic map positions are shown on right- and left-hand side, respectively. Solid lines (in blue and red colour) represent to markers that showed significant associations with traits of interest. Map distances are given in cM and displayed using the MapChart.

### Expression levels of *FT* paralogs explain significant variation in flowering time

To assess whether changes in the expression of different *FT* paralogs could explain the phenotypic variation in flowering time, we examined expression of *FT* paralogs among field-grown plants of all 144 DH lines. Expression levels of all 6 *FT* paralogs displayed significant association with flowering time (p<0.001), with different copies accounting genetic variation for flowering time variably; ranging from 23% (*BnC2.FT*) to *BnC6.FTb* (40%) (Fig. 8A). *FT* homologues; *BnA7.FTb* and *Bn*A7.*FTa* localised near the multiple trait QTL (Supplemental Table S12) could explain 30% and 31% of genetic variation in flowering time. Sequence analyses of the PCR products also confirmed that we are detecting *BnC6.FTb* and *BnA7.FTb* accurately in our assays.

**Fig. 8.**
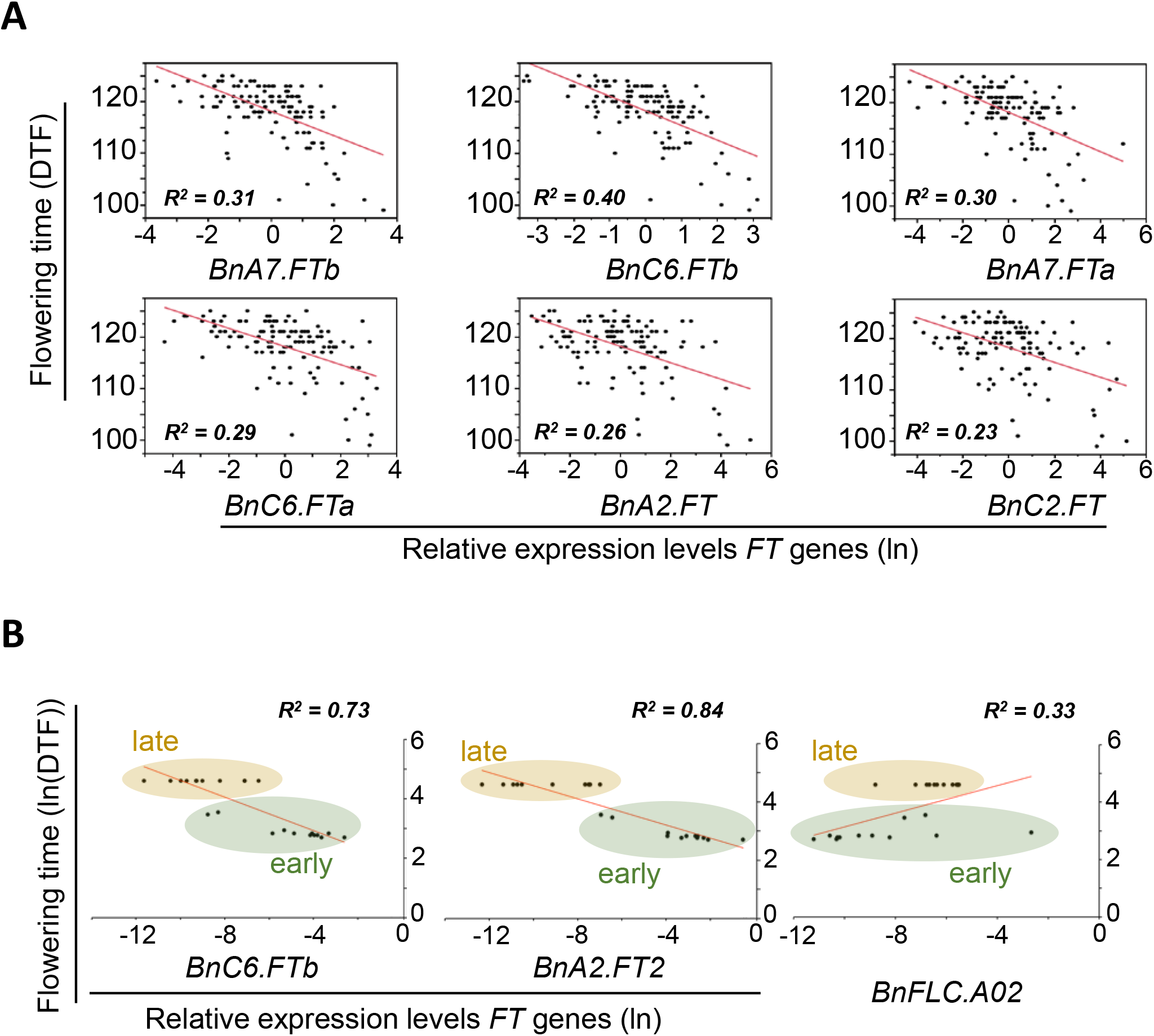
*FT* is a major determinant of flowering time variation and photoperiod gene in canola varieties. A), Expression analysis of different *FT paralogs (BnA2.FTa, BnC2.FT, BnA7.FTa, BnA7.FTb, BnC6.FTa, BnC6.FTb)* on leaves taken from field grown plants of 144 doubled haploid lines of Skipton/Ag-Spectrum//Skipton, and its correlation with flowering time. B), Expression analyses of *FT* genes; *BnC6.FTb (*chromosome C6) and *BnA2.FT* (chromosome A02) and *BnFLC2* on leaves taken from LD grown plants of 24 diverse accessions, representing flowering time diversity in a GWAS panel. The relative expression levels of *FT/and FLC* after normalisation with the reference *UBC9*, is plotted against flowering time.

To further assess whether a similar pattern is also observed among natural variants, we assessed the expression of *BnC6.FTb,* which showed the highest correlation in the DH population and *BnA2.FT2,* which was detected as a QTL in the diversity set of 24 accessions and *BnFLC.A02* in accessions that differed significantly in their flowering time. Consistent with the observations seen in QTL analysis and the expression studies in DH populations, we observed significant differences in *FT* and *FLC* expression that correlated with flowering time among 24 diverse accessions selected on the basis of flowering time diversity (Fig. 8B). Consistent with the timing of sample collection (i.e., just prior to flowering), we detected expression variation in *FT* rather than *FLC* accounted for most of the flowering time variation in these diverse set of 24 accessions. Taken together these data reveal that irrespective of the causal variation, the phenotypic variation is associated with changes in the expression levels of the floral integrator *FT.*

To unravel further *cis* and *trans* acting candidates associated with differential *FT* transcripts expression, we first sought SNPs that affect all *FT* homologues expression levels in diverse canola accessions and then layered this information on the physical map positions of SNPs associated with genetic variation in flowering time and photoperiod response (Supplementary Table S14). We identified a total of 13 SNPs mapped on chromosome A07 and C03, in the vicinity of multiple trait QTLs that we identified in the SAgS population. Candidate genes that were located near significant SNP associations are *FT, ELF4-L2, PRR9, VIN3, BnFLC.C3b (FLC5,* AY036892.1*), FPA, SPA1 and TOE1* (Supplementary Table S11).

### *FT* paralogs exhibit structural sequence variation in *B. napus* accessions

In total, nine *FT* copies were identified in *B. napus* accessions (Supplementary Table 15), including, three putative *FT* copies on chromosomes A01, C02, and C04, (Supplementary Table S15). Sequence analyses showed considerable variation in level of synonymous and non-synonymous SNP variations, Insertion-deletions (InDEL) in promoters, as well as exonic and intronic regions. A total of 310 segregating sites were detected across *FT* paralogs. Our results showed that frequency spectrum of structural variants for *BnA02.FT, BnC02.FT* and *BnC06.FT* conformed to neutral expectations, while *BnC04.FT* and *BnA07.FT* showed non-conformance to neutrality, suggesting evidence of selection. We detected high level of diversity in *FT* paralogs mapped on A07, C04 and C06 chromosomes (Supplementary Table 17). For example, *BnC04.FT* (*BnaC04g14850D*) had 35 SNPs in the genomic sequence, of which the majority of them (21 SNPs) were in intron II. In addition, an 8-bp deletion of the sequence ‘TTCCGGAA’ at coordinates: 12,437,458 to 12,437,465 bp of the *BnC04.FT* was identified in exon-IV among seven accessions; Av-Garnet, BC92157, Skipton, Charlton, BLN3614, ATR-Cobbler, ATR-Gem and in Darmor-*bzh* (reference genotype). As a result, this deletion creates a frameshift mutation that is most likely to alter gene function; the frameshift removes the highly conserved C-terminal domain, removing a large proportion of the PEBP-domain and several substrate-binding sites. Cluster analysis showed that all variants formed a distinct cluster (Figure 9). In the *BnA07.FTb* (BnaA07g33120D) gene, we identified two indel mutations in the coding region that are unlikely to have major effects on protein function. The first is a single nucleotide deletion in exon 3 that is heterozygous with the wild type allele in Australian varieties; Av-Garnet, Skipton, Charlton, BC92156, Marnoo, BLN3614, Ag-Castle, Monty, Maluka, BLN3343-C00402, CB-Telfer, ATR-Gem, Surpass402, ThunderTT, ATR-Mako, Wesroona and Ag-Spectrum (the remaining lines are homozygous wild-type). The deletion results in a frameshift that affects the final 20 amino acids of the encoded peptide, including the 9 amino acids of the PEBP domain. The second InDel is a 3 base-pair mutation in exon 1 (His60-deletion) that is found in all our sequenced lines. These polymorphisms are consisted with the observed QTLs at the vicinity of *FT.*

**Fig. 9.**
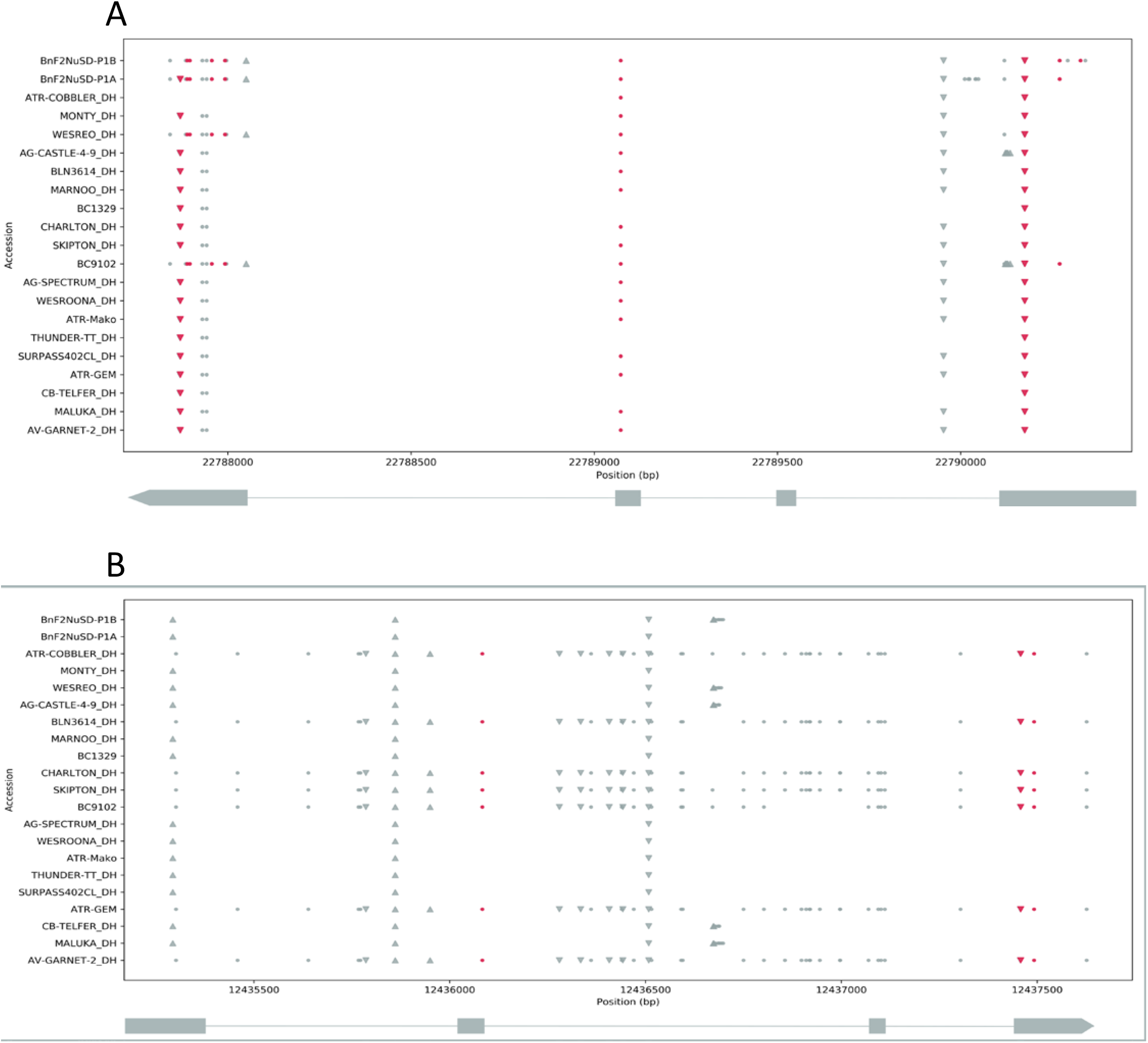
Location of SNPs and indels in FT paralogs *BnaA07g33120D* (A) and *BnaC04g14850D* (B) across the 21 lines representing the GWAS panel and parents. Dots represent SNPs, triangles insertions, and inverted triangles deletions. SNPs and indels shaded in red are non-synonymous. The x-axis shows the scaffold genomic coordinates based on genome version 4.1. The four exon gene model is shown below the plot, with the direction of transcription indicated by the arrow.”

### Structural variation in *FT* promoter region

We further searched CArG box and other motifs for *FLC, SOC1* and *CO* which can potentially bind to repress *FT* expressions (Deng *et al.*, 2011) in introns (especially intron 1) exons and promoter regions. A putative *CO* binding site within Block A: type II = ‘ATTGTGGTGATGAGT’ (Wang et al 2009) was found in both *BnA02.FT* and *BnC02.FT* genes. However, this Type-II block ‘A’ sequence was absent in all *FT* paralogs located on to A07 and C06 chromosomes. There was a single bp deletion in ‘CArG’ box was absent from in introns 1 of *BnA02.FT* and *BnC02.FT* genes. We also found several ‘CACTA’ elements in *B. napus FT* paralogs. For example, in the *BnaC04g14850* gene, a total of four motifs were identified; three were present in introns (2 in Intron 2, antisense direction) and one in sense strand, and one CACTA motif was identified in Exon-IV. In *BnA2.FT*, a total of 834 ‘CACTA’ motifs were identified in promoter, intron 1 and exon II.

In order to determine whether polymorphism in *FT* directly relates to flowering time variation, we performed phylogenetic analysis of 21 accessions representing GWAS panel and parents of mapping populations being used in the Australian Brassica Germplasm Improvement Program. Our results showed that grouping for both spring and winter types based on *FT* paralogs was not that distinct (Fig. 10) suggesting that other key flowering genes such as *FLC* and *FRI* may have contributed to diversification of these morphotypes (Schiessl *et al.*, 2017; Schiessl *et al.*, 2015).

**Fig. 10.**
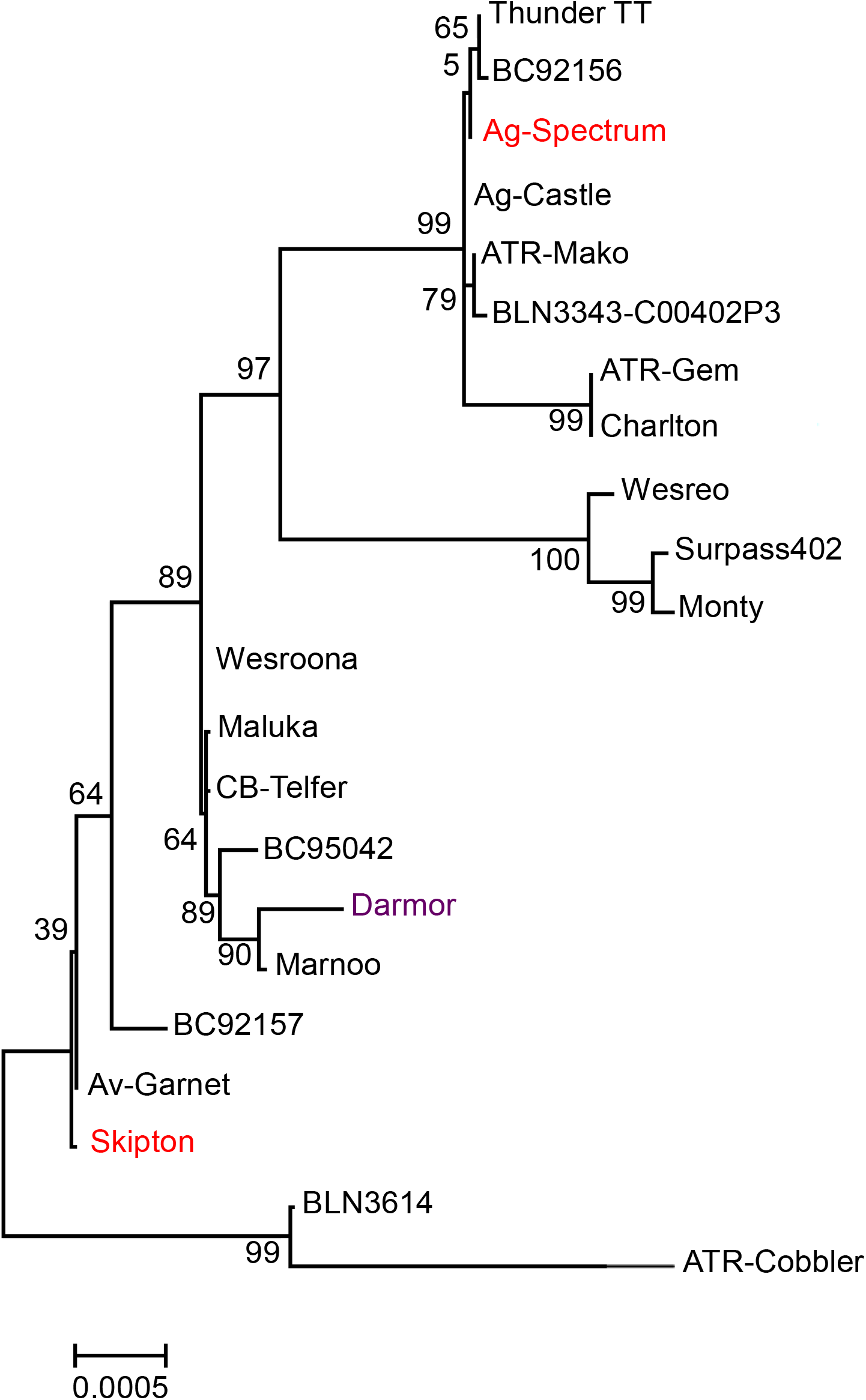
Neighbour-joining tree based on nucleotide variation across all *FT* paralogs among 21 accessions of *Brassica napus* representing GWAS and parental lines (shown in red colour) of a doubled haploid validation population derived from Skipton/Ag-Spectrum//Skipton. Tree was generated in MEGA 6. Nucleotide variation in *FT* genes was also compared with the corresponding *FT* genes in the reference ‘Darmor’ assembly (in purple colour, Supplementary Table x). Number refers to percent bootstrap support for branches with greater than 50% support.

## DISCUSSION

In this study, we explored the genetic architecture of phenotypic diversity in flowering time involved in plant development, adaptation and productivity traits. Our results demonstrate that there is extensive genetically controlled natural variation in flowering time of canola and is due to response to photoperiod (as revealed from LD and SD conditions) and a combination of photoperiod and vernalisation response (Fig. 1). Despite of extended photoperiod at 20°C, several accessions did not flower under CE conditions suggesting that these accessions require vernalisation and flowered when exposed to extended periods of cold temperatures (Raman et al 2016). In order to have a minimum effect of vernalisation on flowering time, all field trials were conducted in the middle of June (instead of April –the main canola growing season in Australia); we identified a highly significant QTL close to *FT* locus on chromosome A02, as identified under CE conditions in LD conditions, suggesting that *FT* is a major candidate for flowering time across different growing conditions (Fig 4). This QTL was also mapped within 80 Kb from QTL for vernalisation response in our previous study (Raman *et al.*, 2016b), suggesting that *FT* integrates signals both from photoperiod and vernalisation pathways in canola. The functional role of *FT* was determined using quantitative RT-PCR using six *FT* paralog specific primers. Our results demonstrated that all paralogs underlie genetic variation in flowering time in canola. For the first time, we showed *FT* expression in a canola population grown under field conditions is significantly associated with variation in flowering time. It was interesting to observe that most of variation in flowering time was explained by A02 locus in GWAS, and A2 and A07 loci near *FT* paralogs in the SAgS DH mapping population (Fig. 6, Supplementary Table S12). However, the maximum correlation (R^2^ = 0.4) was observed for *BnC6.FTb* homologue, followed by *BnA7.FTb* (R^2^ = 0.31), *BnA7.FTa* (R^2^ = 0.30), *BnC6.FTa* (R^2^ = 0.29), *BnA2.FT* (R^2^= 0.26), and *BnC2.FT* (R^2^= 0.23). Higher correlation among different paralogs suggested that different copies can substitute allelic effect on flowering time. Unlike previous studies (Guo *et al.*, 2014; Wang *et al.*, 2012), our results showed that all copies of *FT* are functional. Although all *FT* paralogs except *BnC6.FTa* and *BnC6.FTb* map at the same positions of the closest relative of *FT, TWIN SISTER OF FT* (*TSF*), cloning of six paralogs of *FT* in canola (Wang *et al.*, 2012; Wang *et al.*, 2009) discounted the possibility of *TSF* controlling variation in flowering time which is shown expressed at much lower levels than *FT* (Jang *et al.*, 2009; Michaels *et al.*, 2005; Yamaguchi *et al.*, 2005). No sequence variation was observed in the *FT* paralogs located on chromosomes A01 and C02 among 21 accessions sequenced. Previously, it was reported that these paralogs are evolved may have retained or lost gene function in the polyploid genome of canola (Wang et al. 2009).

We also showed that *FT* has multifaceted role in different plant development, flowering and grain yield, as several QTL were localized in a cluster and *FT* gene expression has shown a good correlation with different traits. However, this relationship was dependent upon G × E interaction (Supplemental Fig. 1). In canola, sequence variation for *BnFLC.A10* appears to underlie QTL for both flowering time as well as root biomass (Fletcher *et al.*, 2016; Fletcher *et al.*, 2015). In addition, flowering time has been implicated in plasticity of water-use efficiency, carbohydrate availability, plant vigour, resistance to diseases and yield (Graf *et al.*, 2010; Kenney *et al.*, 2014; Ni *et al.*, 2009; Wei *et al.*, 2014). We propose that alleles which showed significantly association with flowering time and grain yield in water-limited environments in 2013 and 2014 are of highly relevance even they did not reveal genetic associations in water-unlimited (non-stress environment, 2015 and 2016) and could be exploited in canola breeding programs. Stress environments tend to drive changes in flowering time in Brassica as a result of change in allele frequencies at the flowering time genes (Franks *et al.*, 2016; Franks *et al.*, 2007).

Our findings reveal that the genetic architecture of natural variation in flowering time involves multiple alleles having major effects located near *FT, UPSTREAM OF FLC* and *RAV2* genes on chromosome A02 (Table 1). This is in contrast to genetic variation due to vernalisation requirement which is controlled by multiple alleles across genome (Long *et al.*, 2007; Raman *et al.*, 2016b; Raman *et al.*, 2013). Several SNP markers based on Illumina infinium array were located near the QTL associated with trait variation and known flowering time genes (Bernier and Perilleux, 2005; Dennis and Peacock, 2007; Michaels, 2009). Based on their photoperiodic response, all genotypes could be grouped into photoperiod sensitive, photoperiod insensitive (less sensitive), and non-flowering requiring vernalisation. Clustering of such genotypes based on flowering habit was also supported with our molecular marker based phenetic analysis. The majority of winter types originating from Europe, China and Japan and requiring an extended period of vernalisation to flower seem to be derived from a single cluster (cluster II).

In summary, we have demonstrated through a series of complementary and exploratory analyses based on association tests using genome-wide SNPs, expression QTL and quantitative RT-PCR that the natural variation in flowering time and response to photoperiod revealed in this study is controlled by *FT* and other loci dispersed across the genome, and modulated by the environment. GWA approach delineated genomic regions and provided insights into the genetic architecture of flowering time that control flowering time and its multifaceted role in plant development and productivity traits. Although, some alleles which were identified may not be causative but could be used as selection tools to increase rate of genetic gain in canola improvement programs.

## Acknowledgments

HR thanks Dr. Bev Orchard NSWDPI for advice on controlled environment experiment designs, Dr. Phil Salisbury (DEDJTR and University of Melbourne) for providing seeds of DHC2211 and DHC2261, Mr. Chris Fuller, Mr. Dean McCallum, and Daryl Reardon (NSW Department of Primary Industries) for carrying-out NDVI, and Kristin Verstermark for help with RNA extractions. HR is thankful to Drs Andrzej Killian, Jie Song and Andrew Kowalczyk for supporting KDCompute pipeline for genetic analyses.

This work was supported by grants from the Grains Research and Development Corporation (DAN00117, DAN00208) and NSW Agricultural Genomic Centre, BioFirst Initiative of NSW Government, Australia to HR. SS is supported by an ARC-Australian Post-Doctoral Fellowship (DP110100964) and SB is supported by an ARC-Future Fellowship (FT100100377), Larkins Fellowship and a Linkage Development Scheme from Monash University.

## Authors contributions

HR conceived the research idea and plans; HR, RR, YQ, OO, and IM carried out the phenotypic experiments; HR and RR conducted genotypic analysis; LB, RM, RR and HR analysed data and carried-out trait-marker associations; HR and RR conducted comparative mapping; ASV, SS, HR and SB performed *FT* and *FLC* analyses, HR and DW performed bioinformatics analysis; HR prepared the draft and SB revised it. All authors read/commented the manuscript.

## Supplemental Tables

**Table S1** Accessions used to reveal natural variation in flowering time and photoperiodic response.

**Table S2** Details for phenotyping, experimental designs and QTL analysis

**Table S3** Mean marker density of Illumina SNP markers genotyped in a canola GWAS panel of 368 accessions.

**Table S4** PCR primers used for expression analysis by RT-qPCR (Guo et al 2014)

**Table S5** *Brassica napus* genome BLAT HITs against the *Arabidopsis thaliana FLOWERING LOCUS T* (AT1G65480.1, RSB8/FT/chr1:24331428-24333935) using Darmor reference assembly (http://www.genoscope.cns.fr/blat-server/cgi-bin/colza/webBlat). *FT* paralogs identified in a previous study (Schiessl et al 2014) are also shown for comparision.

**Table S6** (A) Natural variation in flowering time in a GWAS panel of 368 lines of *B. napus* grown under controlled environment cabinets under short day (8 h light and 16 h dark) and long day (16 h light and 8 h dark); (B) Supplemental Table S6b Table. Natural variation in flowering time in a GWAS panel of 300 lines of *B. napus* grown under field conditions. - represents to missing data and (C) Supplemental Table S6C Table: Broad sense heritability of flowering time under controlled and field condition among canola accessions.

**Table S**7 Marker LD across *B. napus* genome.

**Table S8** Familial relationships between pairs of accessions used for GWAS.

**Table S9** Marker trait association identified for flowering time and photoperiodic response in a GWAS panel of canola. Response to photoperiod was assessed under controlled environment conditions, LD: Long day conditions (16 h light, 8 hr dark at 20 degree); SD (8 h light, 16 h dark at 20 degree). Flowering time was also evaluated under field conditions at three sites: Wagga Wagga (irrrigation, NSW, Australia), Wagga Wagga (lateral site) and Condobolin (rainfed site, NSW, Australia) Days to flowering was used for GWAS analysis using GAPIT program in R and Golden Helix (SVS, with and without principal component analysis).

**Table S10** Distribution of significant marker associations for flowering time and photoperiod response, evaluated under controlled environment cabinets and field conditions (three sites) in a GWAS panel of canola

**Table S11** Candidate gene associated with flowering time and photoperiodic response in the GWAS and DH population.

**Table S12** Significant QTL associated with flowering time and grain yield identified in a doubled haploid population derived from a single BC_1_F_1_ from the Skipton/Ag-Spectrum//Skipton population grown in four environments, at Euberta (2013) and Wagga Wagga (2014, 2015 and 2016). QTL in bold are repeatedly detected across environments/traits. QTL in bold and italics are multi-trait QTL (pleiotropic).

**Table S13** Genetic correlation between different traits measured in the doubled haploid population from Skipton/Ag-Spectrum//Skipton across environments.

**Table S14** Genome-wide association analysis (eQTL) showing statistical association between Illumina SNP markers and expression data of *BnC6.FT* gene in 300 accessions of *B. napus*. Linear marker regression analysis was performed in the SVS package (Golden Helix).

**Table S15** Summary of structural and polymorphic variation identified among 21 *B. napus* accessions representing GWAS and validation population used in this study. Numbers in table represent counts of unique variants observed across the 21 accessions. Abbreviations: SNV: structural nucleotide variant, InDel: Insertion-deletion, S = Number of segregating sites, ps = S/n, Θ = ps/a1, π = nucleotide diversity, and D is the Tajima test statistic (Tajima, 1989).

**Table S16** Gene structures of different *FT* paralogs identified in the resequence data from 21 accessions of *B. napus* (test samples). Exon/intron genomic coordinates of the *B. napus* reference cultivar are based on the current gene models (annotation version 5). Numbers in the table represent lengths in base-pairs. Exon/intron length variation in the 21 accessions (in bold) is only counted for InDels that are homozygous.

---

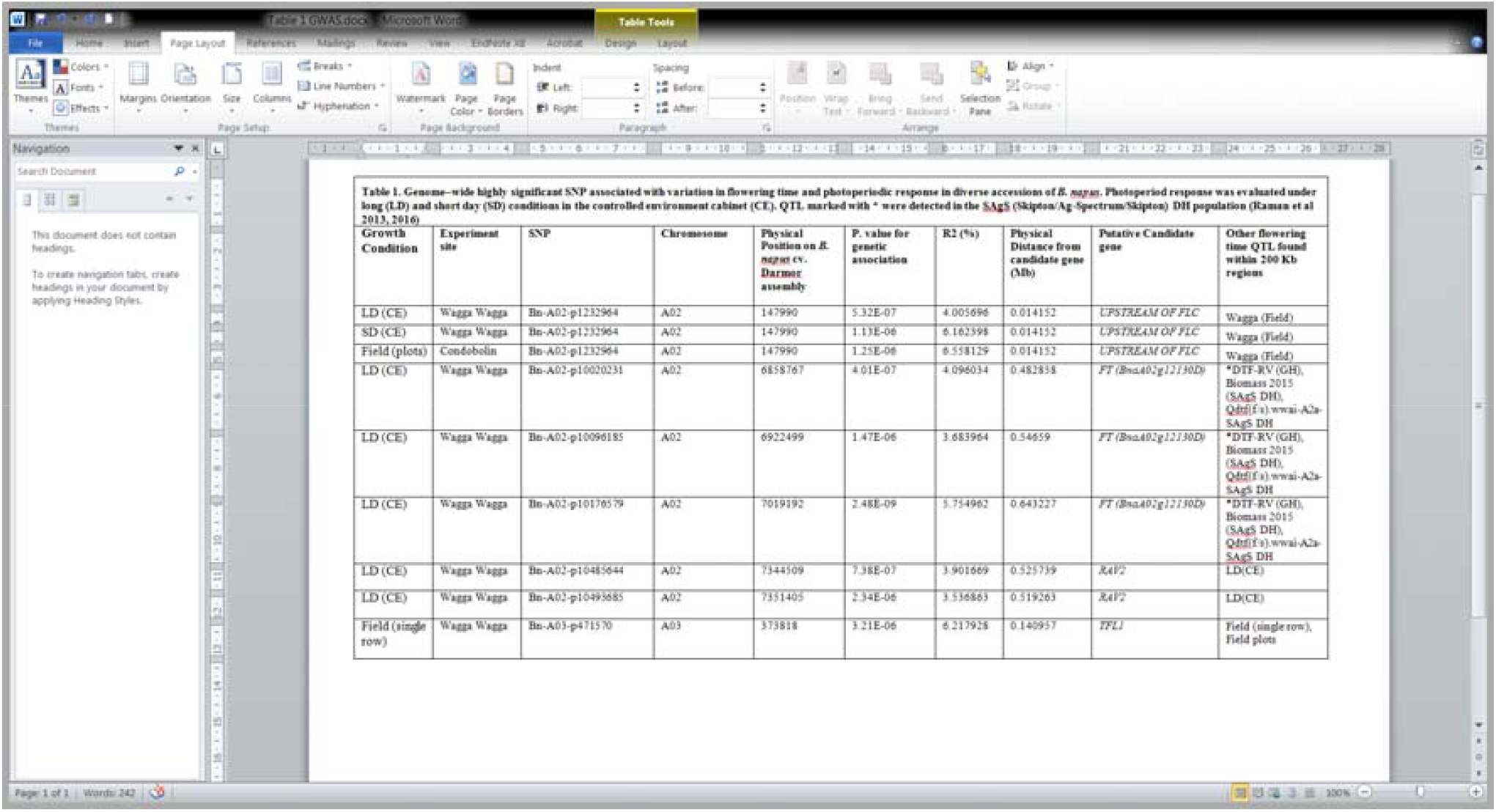

