## Supplementary Table2 for "GWAS hints at pleiotropic roles for *FLOWERING LOCUS T* in flowering time and yield-related traits in canola"

**Supplemental Table S2.**

**Layout of Wagga Wagga Trial-2017 (layout)**

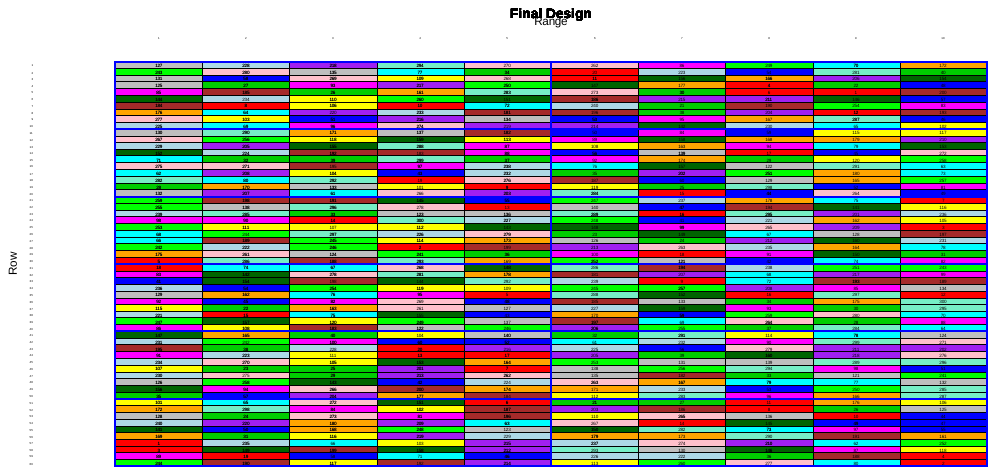

**Layout of Condobolin Trial-2017**

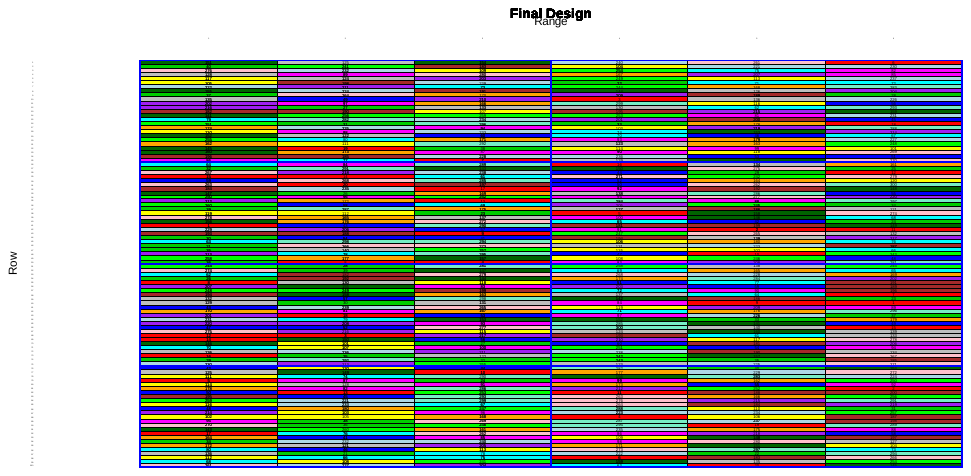

**Layout of Wagga Row Trial-2017**

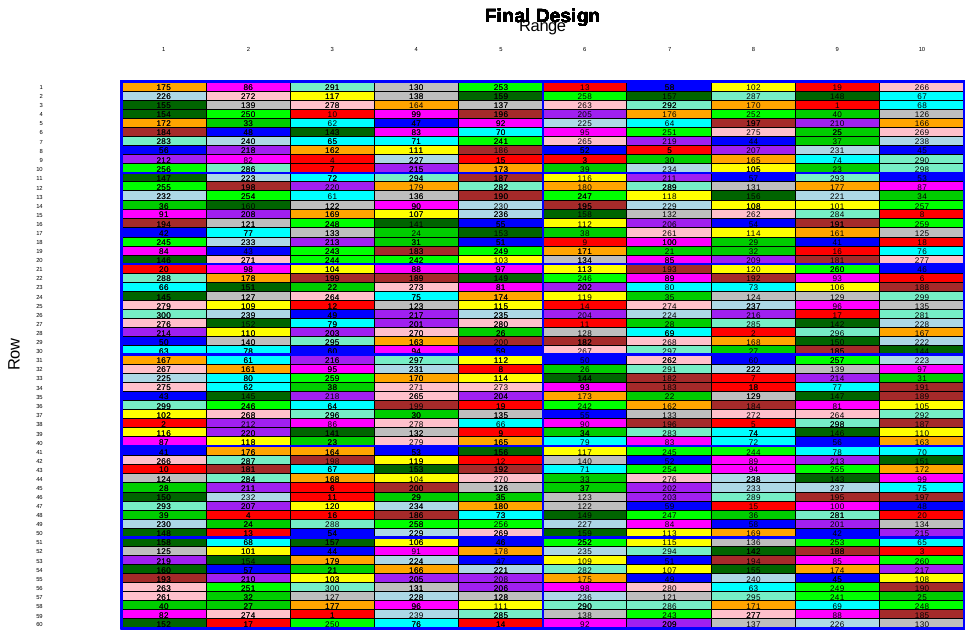

**Experiment design for investigating the role of *FT* and *FLC* in flowering time under controlled cabinet**

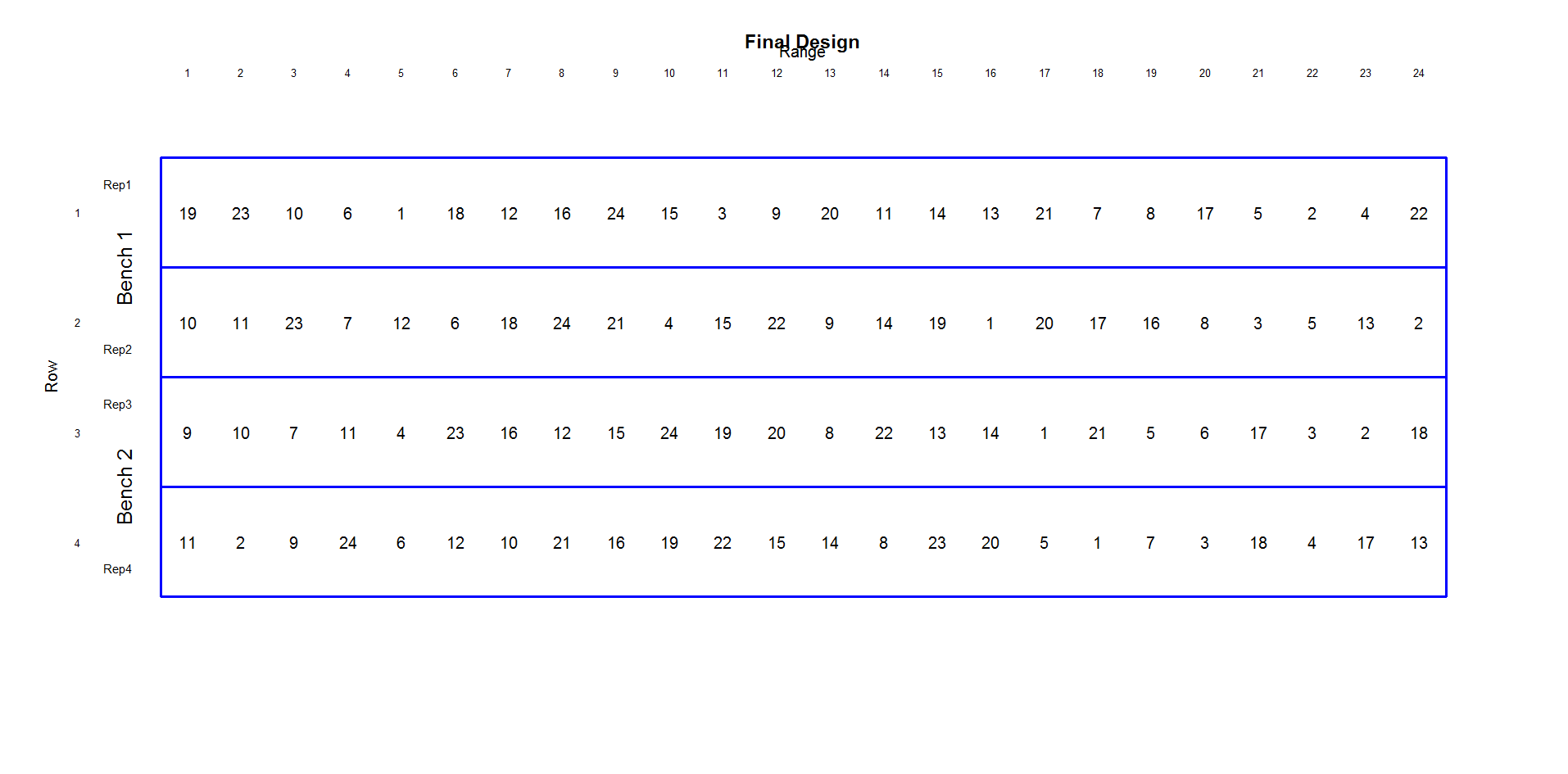

**Details of diverse accessions representing GWAS panel used for FT and *FLC* expression analyses**

Accession name	Accession Number

| ZHONGYOU-ZA-NO8_DH | 15352 |
| --- | --- |
| Beluga | 51490 |
| GSL1-1_DH | 52505 |
| ZY014-8_DH | 52570 |
| RANGI-501-12_DH | 52586 |
| PRIMOR-1_DH | 52611 |
| NORIN-19-469973-11_DH | 52612 |
| AZUMA-13_DH | 52633 |
| TOWER | 95048 |
| ERGLU | 95050 |
| WESBELL_DH | 52387 |
| TATYOON_DH | 52390 |
| CB-TRILOGY_DH | 52412 |
| ROTTNEST-TTC_DH | 52422 |
| SARDI524TT-11_DH | 52555 |
| AGT346-11_DH | 52659 |
| ATR-SNAPPER_DH | 94504 |
| CB-JARDEE-HT-16_DH | 94551 |
| Rapid cycling | 95194 |
| Columbus | 50557 |
| Darmor | 51831 |
| CB-Telfer | 52430 |
| Skipton | 52649 |
| Ag-Spectrum | 52374 |
