## Supplementary Table 12 for "GWAS hints at pleiotropic roles for *FLOWERING LOCUS T* in flowering time and yield-related traits in canola"

**Supplemental Table S12a.** Heritability of different phenotypic measurements in a DH population from Skipton/Ag-Spectrum//Skipton

| Trait | Phenotyping year | H^2^ (%) |
| --- | --- | --- |
| Flowering time | 2013 | 74 |
|  | 2014 | 98 |
|  | 2015 | 95.55 |
|  | 2016 | 97.07 |
| Plant emergence | 2016 | 44.28 |
| Biomass | 2015 | 65.92 |
|  | 2016 | 56.73 |
| Plant height | 2015 | 70.12 |
|  | 2016 | 83.8 |
| NDVI (2^nd^ July) | 2015 | 29.24 |
| NDVI (8^th^ July) | 2015 | 87.38 |
| NDVI (17^th^ July) | 2015 | 89.25 |
| NDVI (29^th^ July) | 2015 | 92.48 |
| NDVI (1) | 2016 | 79.75 |
| NDVI (2) | 2016 | 83.31 |
| NDVI (3) | 2016 | 88.17 |
| NDVI (4) | 2016 | 88.04 |
| NDVI (5) | 2016 | 77.71 |
| Grain yield | 2013 | 78 |
|  | 2014 | 99 |
|  | 2015 | 90.87 |
|  | 2016 | 95.98 |

**Supplemental Table S12b: Significant QTL associated with flowering time and grain yield identified in a doubled haploid population population derived from a single BC_1_F_1_ from the Skipton/Ag-Spectrum//Skipton population grown in four environments, at Euberta, NSW, Australia(2013) and Wagga Wagga, NSW, Australia(2014, 2015 and 2016). QTL in bold are repeatedly detected across environments/traits. QTL in bold and italics are multi-trait QTL (pleiotropic).* Physical map positions are based on the consensus map of *B. napus*.**

| Trait | Phenotyping year | Marker locus | Chromosome | Distance (cM) | Allelic Effect | LOD | R2 | Physical map position on Darmor genome | Consistent QTL |
| --- | --- | --- | --- | --- | --- | --- | --- | --- | --- |
| Grain yield | 2014 | 3075958 | A01 | 44.53 | 0.09 | 1.8 | 3 | 20301620 |  |
| Biomass | 2016 | 3185901 | A01 | 55.39 | 0.48 | 2.18 | 2.43 | 22569015 |  |
| **Days to first flower** | **2015** | **3107075** | **A02** | **2.33** | **-1.61** | **7.57** | **9.81** | ***2288953** | **1** |
| **Days to last flower** | **2015** | **3107075** | **A02** | **2.33** | **-0.88** | **3.63** | **4.42** | ***2288953** | **1** |
| **Days to last flower** | **2016** | **3107075** | **A02** | **2.33** | **-1.28** | **3.05** | **3.61** | ***2288953** | **1** |
| Days to first flower | 2016 | 3082031 | A02 | 8.53 | -1.51 | 4.78 | 6 | 217384 |  |
| Height | 2015 | 3145775 | A02 | 9.31 | -2 | 4.58 | 5.72 | 218367 |  |
| Days to first flower | 2014 | 3168973 | A02 | 10.08 | -1.4 | 2 | 3.6 | 262474 |  |
| Plant emergence | 2016 | 3088452 | A02 | 26.68 | 2.02 | 3.77 | 4.61 | 2579820 |  |
| Biomass | 2015 | 3081578 | A02 | 37.55 | -0.72 | 3.9 | 4.79 | 4012023 |  |
| Biomass | 2015 | 3086516 | A03 | 0.77 | -0.52 | 2.37 | 2.69 | 13568216 |  |
| Grain yield | 2014 | 3113944 | A04 | 38.8 | -0.1 | 2.6 | 5 | 14743951 |  |
| ***Grain yield*** | ***2014*** | ***4109113_65:A>G*** | ***A06*** | ***3.88*** | ***0.1*** | ***3.9*** | ***8.9*** | ***927363*** | ***2*** |
| ***NDVI*** | ***2016*** | ***4109113_65:A>G*** | ***A06*** | ***3.88*** | ***-0.01*** | ***2.84*** | ***3.33*** | ***927363*** | ***2*** |
| Grain yield | 2013 | 3155700 | A06 | 10.09 | 0.4 | 2.5 | 5 | 1792355 |  |
| Biomass | 2014 | 3100269 | A06 | 34.18 | -44.80 | 2.03 | 5.13 | 2713555 |  |
| Height | 2015 | 3136779 | A06 | 37.25 | -1.88 | 3.82 | 4.67 | 3409201 |  |
| NDVI | 2016 | 3110950 | A06 | 43.46 | -0.01 | 5.26 | 6.66 | 4902896 |  |
| Grain yield | 2013 | 4120436 | A06 | 44.23 | -0.4 | 2.4 | 4.5 | 4982032 |  |
| Grain yield | 2016 | 3154085* | A06 | 53.58 | -0.1 | 3.21 | 3.84 | 6409628 |  |
| Grain yield | 2015 | 4113040 | A07 | 10.86 | -0.07 | 3.32 | 3.99 | *11765705 |  |
| NDVI | 2015 | 4106860 | A07 | 12.41 | -0.02 | 2.57 | 2.97 | 14817135 |  |
| **Height** | **2015** | **4106995** | **A07** | **60.19** | **-3.64** | **12.5** | **16.25** | **22280076** | **3** |
| **Height** | **2016** | **4106995** | **A07** | **60.19** | **-2.13** | **3.43** | **4.14** | **22280076** | **3** |
| Biomass | 2016 | 3157763 | A07 | 61.74 | -1.11 | 9.35 | 12.17 | 22661268 |  |
| ***Grain yield*** | ***2016*** | ***3110489*** | ***A07*** | ***64.85*** | ***-0.1*** | ***3.18*** | ***3.79*** | ***21980170*** | ***4*** |
| ***Grain yield*** | ***2013*** | ***3110489*** | ***A07*** | ***64.85*** | ***-0.5*** | ***3.3*** | ***6.9*** | ***21980170*** | ***4*** |
| ***Days to first flower*** | ***2014*** | ***3110489*** | ***A07*** | ***64.85*** | ***2.3*** | ***4.7*** | ***10.3*** | ***21980170*** | ***4*** |
| ***Biomass*** | ***2015*** | ***3075574*** | ***A07*** | ***65.62*** | ***-1.73*** | ***16*** | ***22.56*** | ***23528818*** | ***5*** |
| ***Biomass*** | ***2014*** | ***3075574*** | ***A07*** | ***65.62*** | ***-85.41*** | ***5.60*** | ***16.03*** | ***23528818*** | ***5*** |
| ***Plant emergence*** | ***2016*** | ***3075574*** | ***A07*** | ***65.62*** | ***1.82*** | ***3.33*** | ***4.01*** | ***23528818*** | ***5*** |
| Days to first flower | 2013 | 4108720 | A07 | 66.4 | 0.3 | 1.8 | 5.9 | 23611343 |  |
| Grain yield | 2014 | 4116554 | A07 | 67.95 | -0.1 | 4.4 | 10.3 | 22902708 |  |
| Biomass | 2015 | 3082330_21:A>T | A10 | 19.56 | -0.8 | 4.88 | 6.14 | 10552105 |  |
| Grain yield | 2014 | 3191929 | C01 | 0 | -0.1 | 3.9 | 9.1 | 1024749 |  |
| Days to first flower | 2013 | 3172867 | C01 | 10.09 | 0.3 | 2 | 6.6 | 2292278 |  |
| Grain yield | 2014 | 3099995 | C02 | 0.77 | -0.1 | 1.5 | 2.5 | 3780953 |  |
| Days to first flower | 2016 | 3133962 | C02 | 3.87 | 1.11 | 2.42 | 2.76 | 26765802 |  |
| Biomass | 2014 | 3114751 | C02 | 1.71 | 47.01 | 2.09 | 5.29 | 6916101 |  |
| Days to last flower | 2016 | 3084832 | C02 | 6.2 | 1.38 | 2.73 | 3.18 | 4686995 |  |
| NDVI | 2015 | 3100600_29:C>G | C02 | 40.07 | 0.01 | 2.22 | 2.49 | 47080045 |  |
| **Days to first flower** | **2015** | **3097882** | **C02** | **65.07** | **0.82** | **2.43** | **2.77** | **45026595** | **6** |
| **Days to first flower** | **2014** | **3097882** | **C02** | **65.07** | **1.5** | **2.5** | **6** | **45026595** | **6** |
| ***Grain yield*** | ***2013*** | ***3077306*** | ***C03*** | ***1.55*** | ***-0.4*** | ***2*** | ***4.9*** | ***735747*** | ***7*** |
| ***Grain yield*** | ***2015*** | ***3077306*** | ***C03*** | ***1.55*** | ***-0.1*** | ***4.19*** | ***5.19*** | ***735747*** | ***7*** |
| ***Days to last flower*** | ***2016*** | ***3077306*** | ***C03*** | ***1.55*** | ***-1.55*** | ***3.73*** | ***4.56*** | ***735747*** | ***7*** |
| ***Grain yield*** | ***2016*** | ***3077306*** | ***C03*** | ***1.55*** | ***-0.25*** | ***11.15*** | ***14.53*** | ***735747*** | ***7*** |
| ***Height*** | ***2016*** | ***3077306*** | ***C03*** | ***1.55*** | ***-2.64*** | ***4.66*** | ***5.84*** | ****735747*** | ***7*** |
| Grain yield | 2014 | 3081698 | C03 | 2.32 | -0.2 | 6.3 | 19 | 3661975 |  |
| Days to first flower | 2013 | 4119165 | C03 | 6.2 | -0.5 | 2.8 | 9.8 | 2605790 |  |
| Height | 2015 | 4120475 | C03 | 17.85 | -1.69 | 2.87 | 3.37 | 4280194 |  |
| **Days to first flower** | **2014** | **4119157** | **C03** | **26.39** | **-2.4** | **4.5** | **9.5** | **5308348** | **8** |
| **Days to first flower** | **2015** | **4119157** | **C03** | **26.39** | **-1.43** | **5.48** | **6.97** | **5308348** | **8** |
| **Days to first flower** | **2016** | **4119157** | **C03** | **26.39** | **-1.44** | **4.16** | **5.14** | **5308348** | **8** |
| NDVI | 2016 | 4119194 | C03 | 32.62 | -0.01 | 2.32 | 2.62 | 47688568 |  |
| Days to last flower | 2015 | 3086134 | C03 | 35.72 | -0.88 | 3.34 | 4.02 | 11986941 |  |
| Grain yield | 2014 | 4119821 | C03 | 36.5 | 0.2 | 6.4 | 17.1 | 6320326 |  |
| Days to last flower | 2016 | 3150474 | C03 | 38.05 | -1.25 | 2.35 | 2.66 | 8072879 |  |
| Grain yield | 2016 | 3113785 | C03 | 42.7 | 0.14 | 4.1 | 5.06 | 13114289 |  |
| **Grain yield** | **2015** | **3090240** | **C03** | **58.99** | **0.07** | **3.8** | **4.65** | **15746555** | **9** |
| **Grain yield** | **2013** | **3090240** | **C03** | **58.99** | **0.3** | **2** | **3.6** | **15746555** | **9** |
| **Grain yield** | **2014** | **3090240** | **C03** | **58.99** | **0.1** | **1.6** | **3.9** | **15746555** | **9** |
| Days to first flower | 2015 | 3080309 | C04 | 2.33 | 1.28 | 3.07 | 3.65 | 6570070 |  |
| Height | 2016 | 3114172 | C04 | 3.1 | 1.67 | 2.08 | 2.3 | 6834294 |  |
| Days to first flower | 2014 | 3091447 | C04 | 11.65 | 2.2 | 2.3 | 6.2 | 62149093 |  |
| **Days to first flower** | **2016** | **3081582** | **C04** | **12.42** | **1.29** | **2.49** | **2.85** | **14611482** | **10** |
| **Days to last flower** | **2016** | **3081582** | **C04** | **12.42** | **1.71** | **3.4** | **4.1** | **14611482** | **10** |
| Days to last flower | 2015 | 3075169 | C04 | 14.75 | 0.89 | 2.09 | 2.31 | 121513392 |  |
| NDVI | 2015 | 3146589 | C04 | 26.39 | -0.02 | 2.67 | 3.09 | Unknown |  |
| Grain yield | 2016 | 3093112 | C04 | 67.02 | 0.07 | 2.05 | 2.26 | 51836170 |  |
| Plant emergence | 2016 | 3105078 | C05 | 35.89 | 1.35 | 2.12 | 2.35 | *4806733 |  |
| **Days to first flower** | **2013** | **4112095** | **C05** | **61.5** | **-0.3** | **2.1** | **6.8** | **Unknown** | **11** |
| **Days to first flower** | **2014** | **4112095** | **C05** | **61.5** | **-1.2** | **1.6** | **2.9** | **Unknown** | **11** |
| Days to first flower | 2014 | 3114192 | C06 | 7.78 | 1.4 | 2.1 | 3.8 | 29379493 |  |
| Days to first flower | 2013 | 4111646 | C06 | 21.74 | -0.3 | 1.9 | 6 | 22133410 |  |
| NDVI | 2016 | 3077957_37:T>G | C06 | 56.02 | 0.01 | 2.04 | 2.25 | 3702996 |  |
| Biomass | 2014 | 3075116 | C09 | 1.69 | -47.93 | 2.25 | 5.66 | 268170 |  |
| Height | 2015 | 4120960 | C09 | 5.43 | -1.53 | 3.17 | 3.78 | 95496 |  |
| Grain yield | 2015 | 3123841 | C09 | 132.36 | -0.08 | 4.58 | 5.73 | 46484334 |  |
| Grain yield | 2013 | 3080127 | C09 | 143.23 | -0.4 | 2.4 | 4.6 | 48475846 |  |
| NDVI | 2015 | 3083679 | C09 | 144 | -0.02 | 5.27 | 6.68 | Unknown |  |
| NDVI | 2016 | 4118386 | C09 | 145.55 | -0.01 | 4.64 | 5.81 | Unknown |  |
