## Supplementary Figures for "GWAS hints at pleiotropic roles for *FLOWERING LOCUS T* in flowering time and yield-related traits in canola"

Supplemental Figure 1. Canola genotypes showing GX E interactions when grown under LD and SD conditions in controlled environment cabinet. Mean flowering time is estimated in days. Details of varieties shown here represented to BC accessions (Supplemental Table S1).

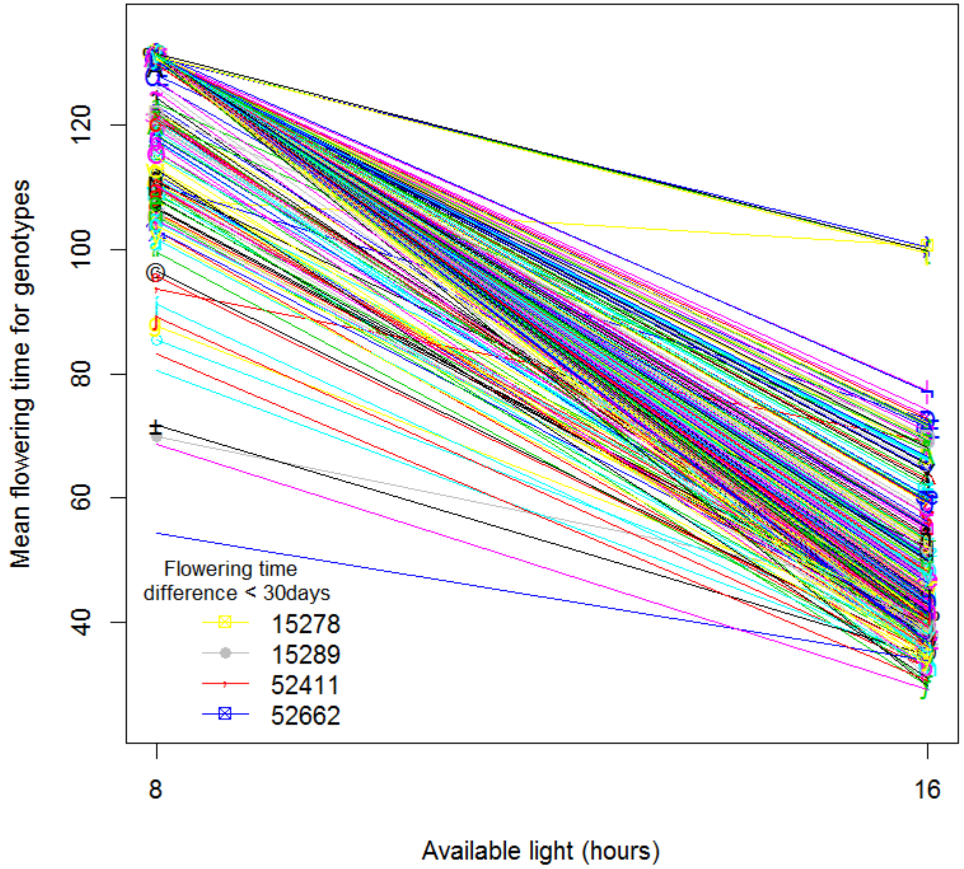

Supplemental Figure 2. Genome-wide distribution (A) and density (B) of single nucleotide polymorphisms, in a genome wide association diversity panel of 368 *Brassica napus* accessions. Regions that are rich and poor SNP density are shown in dark and white horizontal bars, respectively. The number of SNP markers anchoring on different chromosomes (A1-A10 and C1-C9) of the physical map of the *B. napus* genome is given on the x-axis.

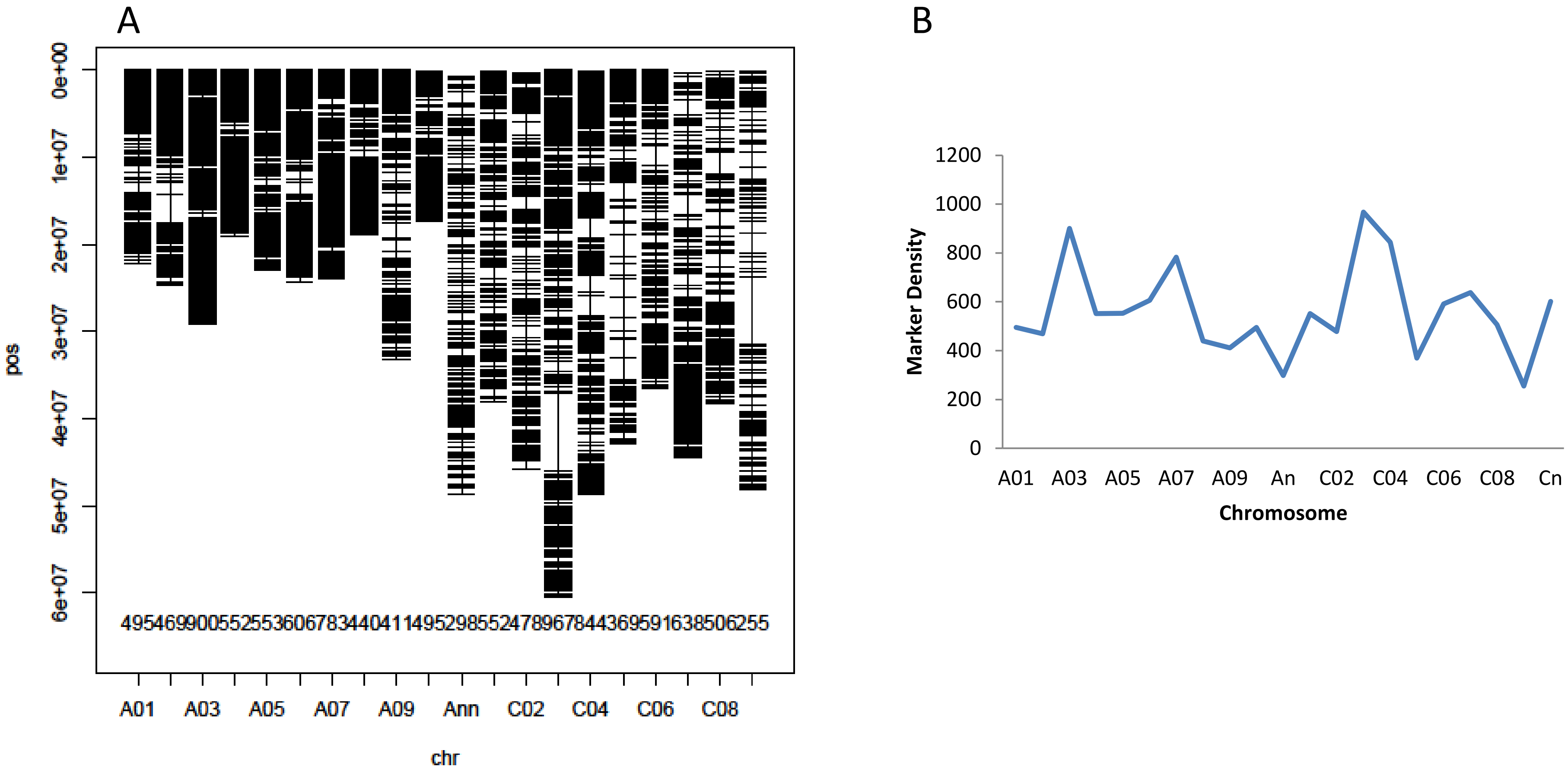

Supplemental Figure 3: Genetic diversity and population structure in a GWAS panel of 368 *Brassica napus* accessions. Three clusters designated as I, II and III represent predominantly Chinese, European, and Australian accessions, respectively. Details of accessions are given in Supplemental Table S1.

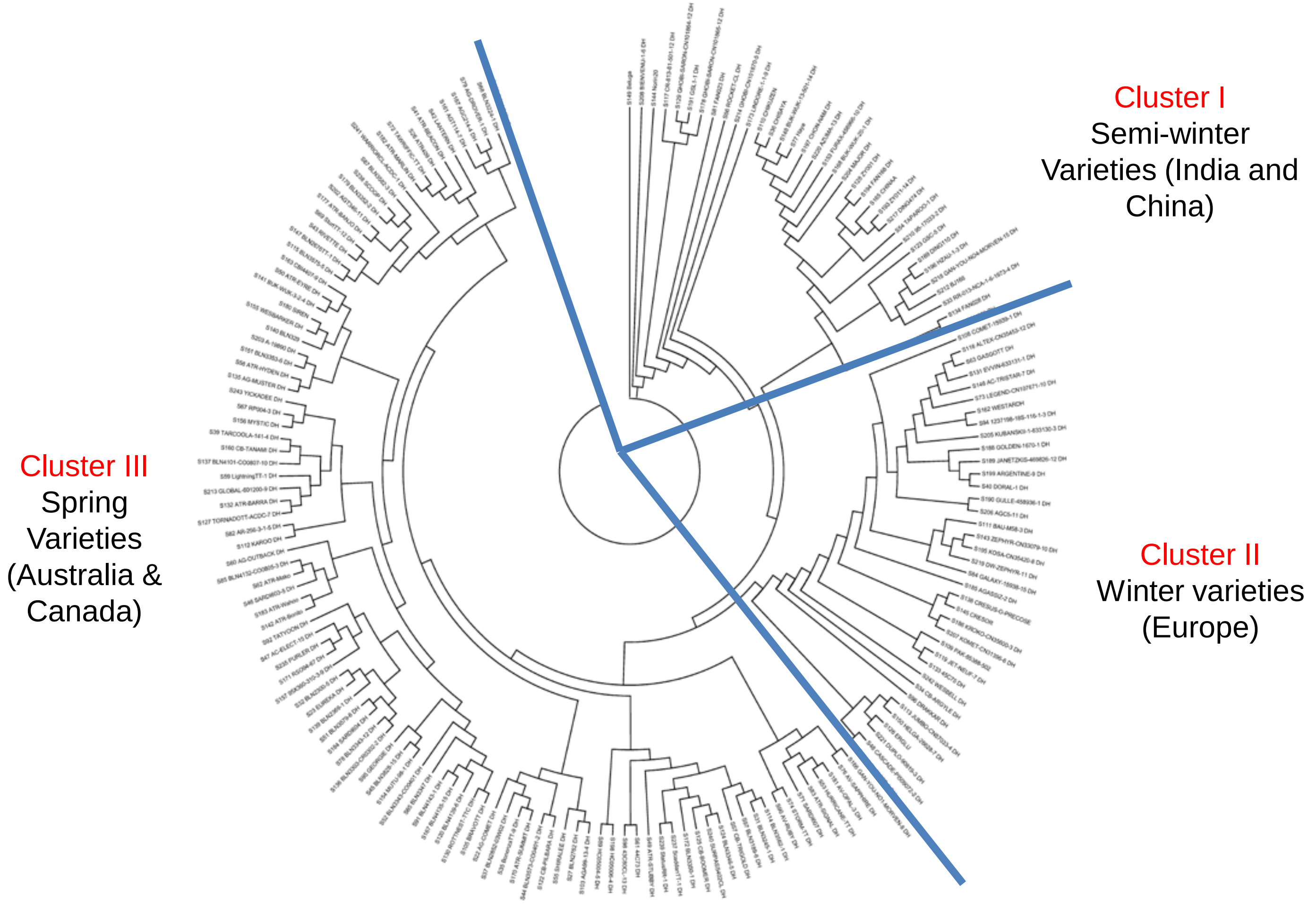

Supplemental Figure 4: Principal components (PC1 and PC2) analysis showing population structure in a GWAS diversity panel of 368 *B. napus* accessions. Three major clusters designated as I, II, and III, consistent with the cluster analysis (Supplemental Figure 2).

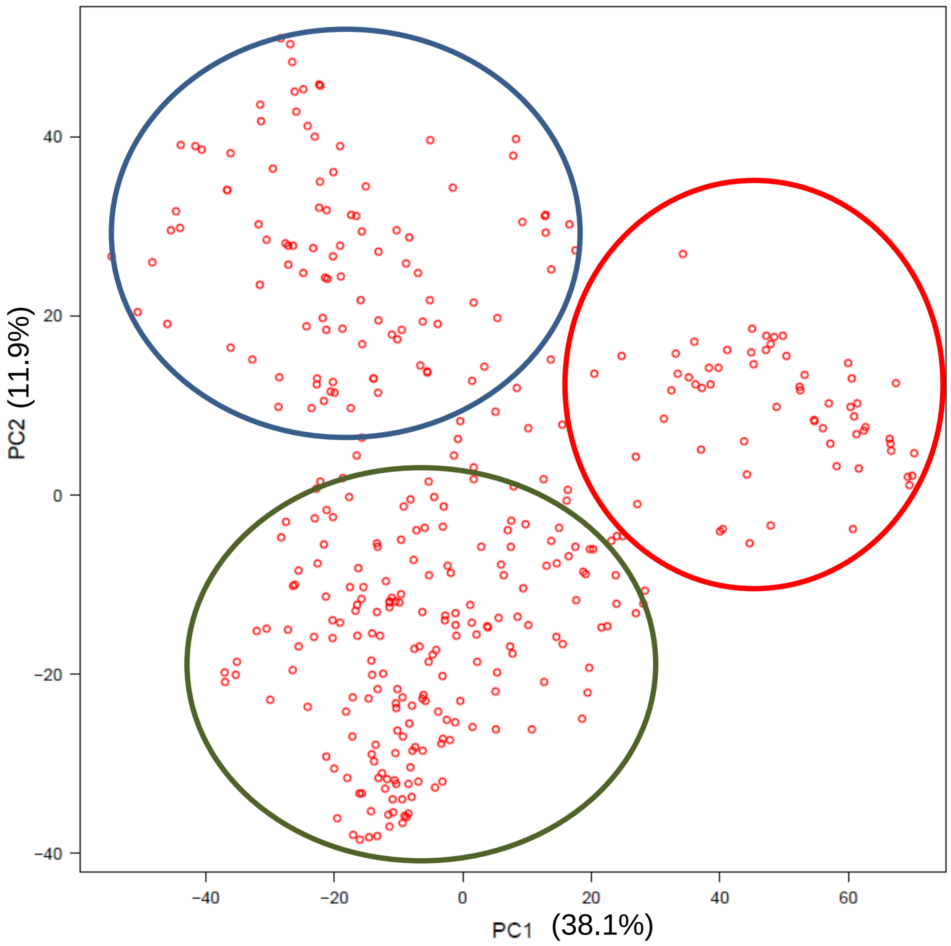

Supplemental Figure 5. The average linkage disequilibrium (LD) decays ($r^2$) approach 0.02 when distance between SNPs was approximately 200 Kb. Distance in bp is shown on X-axis.

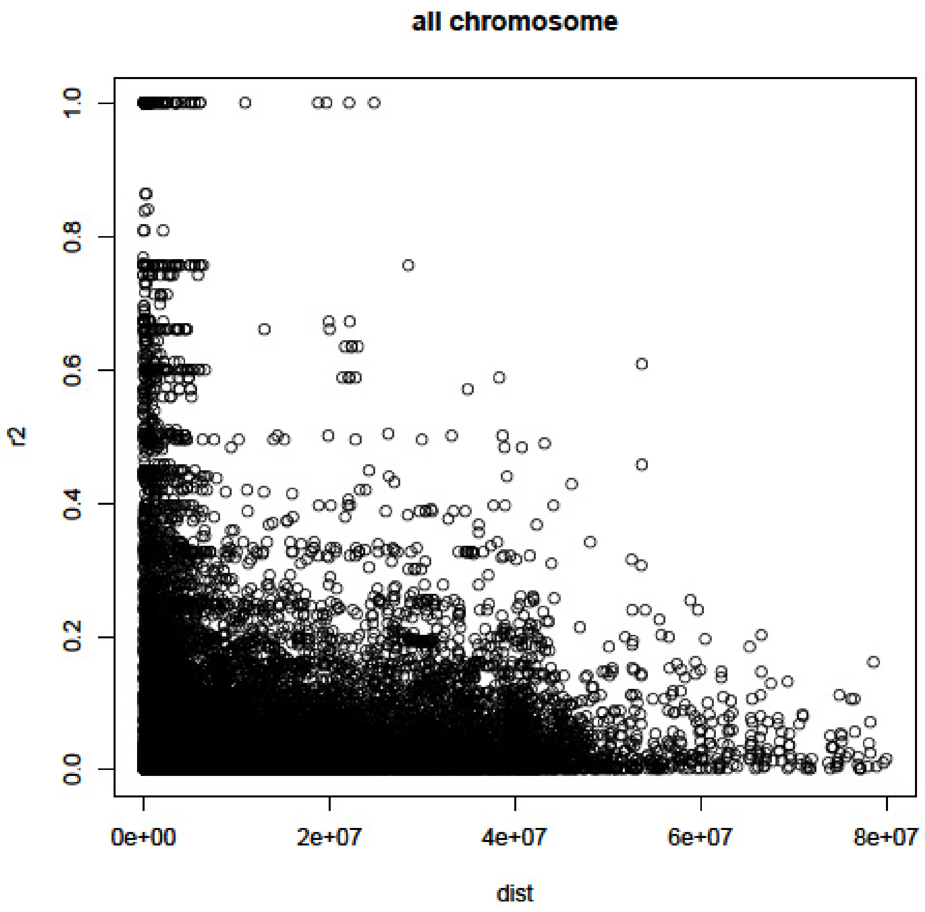

Supplemental Figure 6 (A). Frequency distribution of shoot biomass in a SAgS DH population phenotyped across 2014-2016 growing environments.

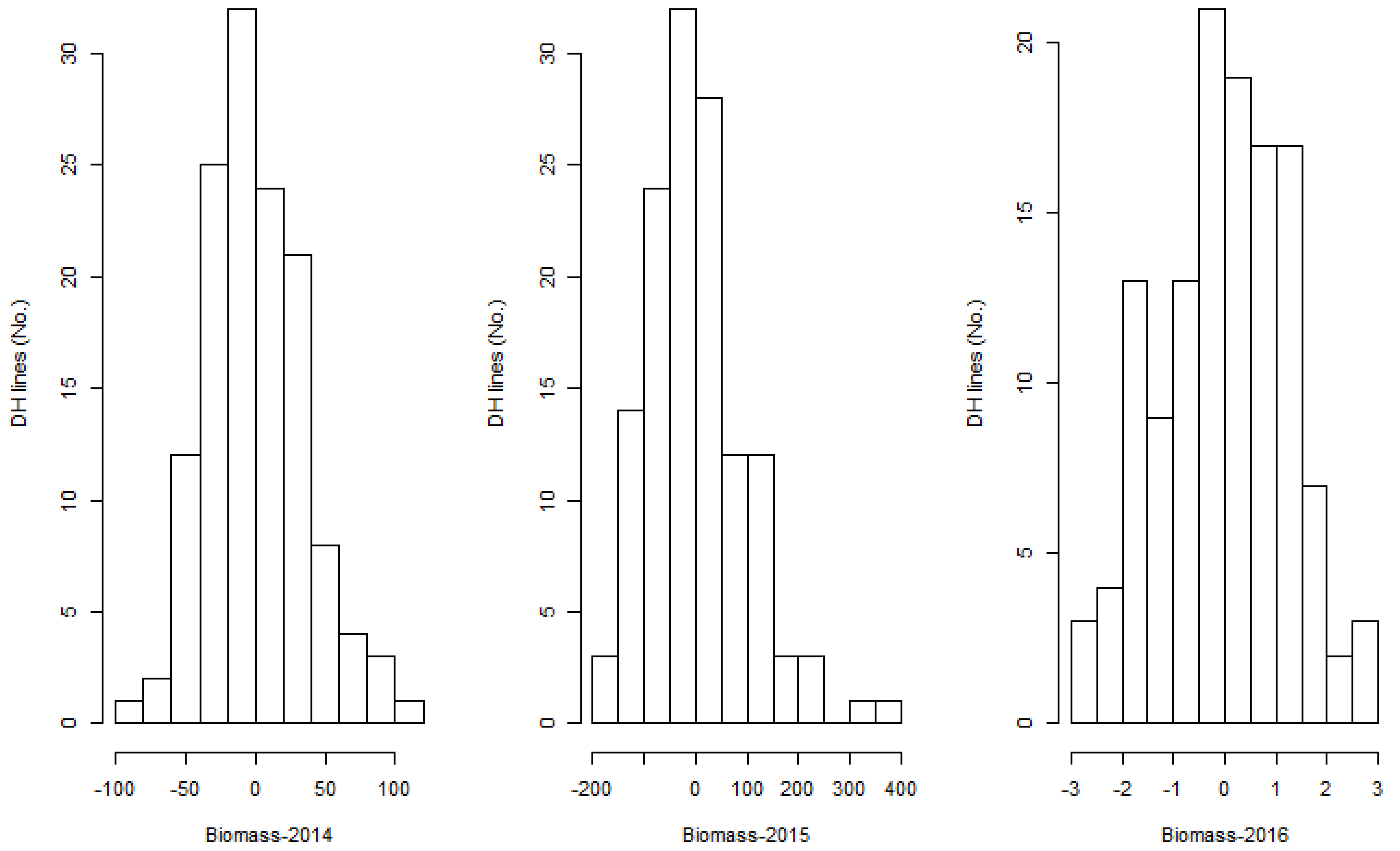

Supplemental Figure 6 (B). Frequency distribution of fractional ground cover, measured as NSVI with a hand-held GreenSeeker machine, in a SAgS DH population phenotyped across 2015-2016 growing environments).

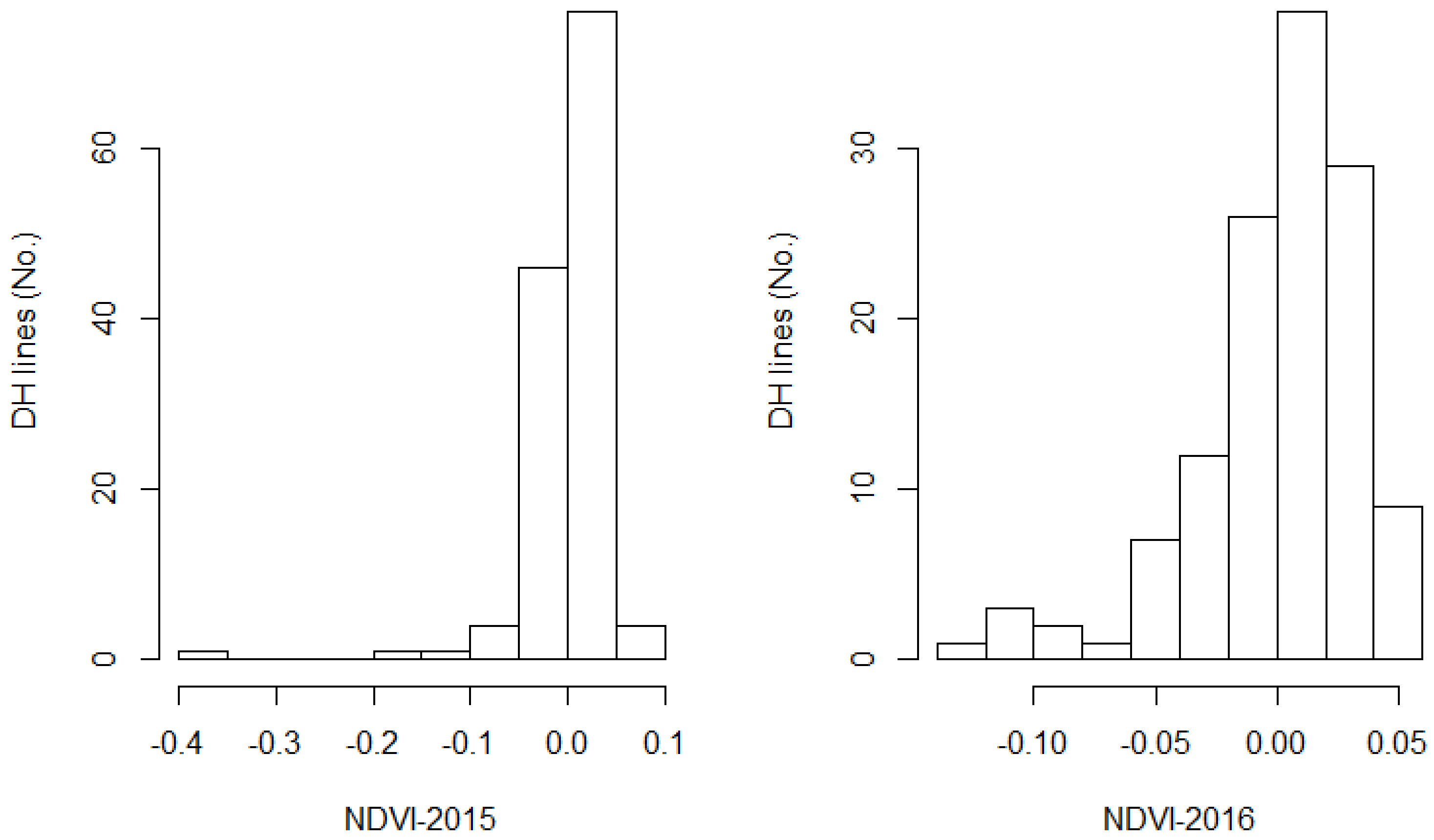

Supplemental Figure 6 (C). Frequency distribution of days to flower in a SAgS DH population phenotyped across four environments (2013-2016). Phenotypic data of 2013 and 2014 was published previously (Raman et al 2016).

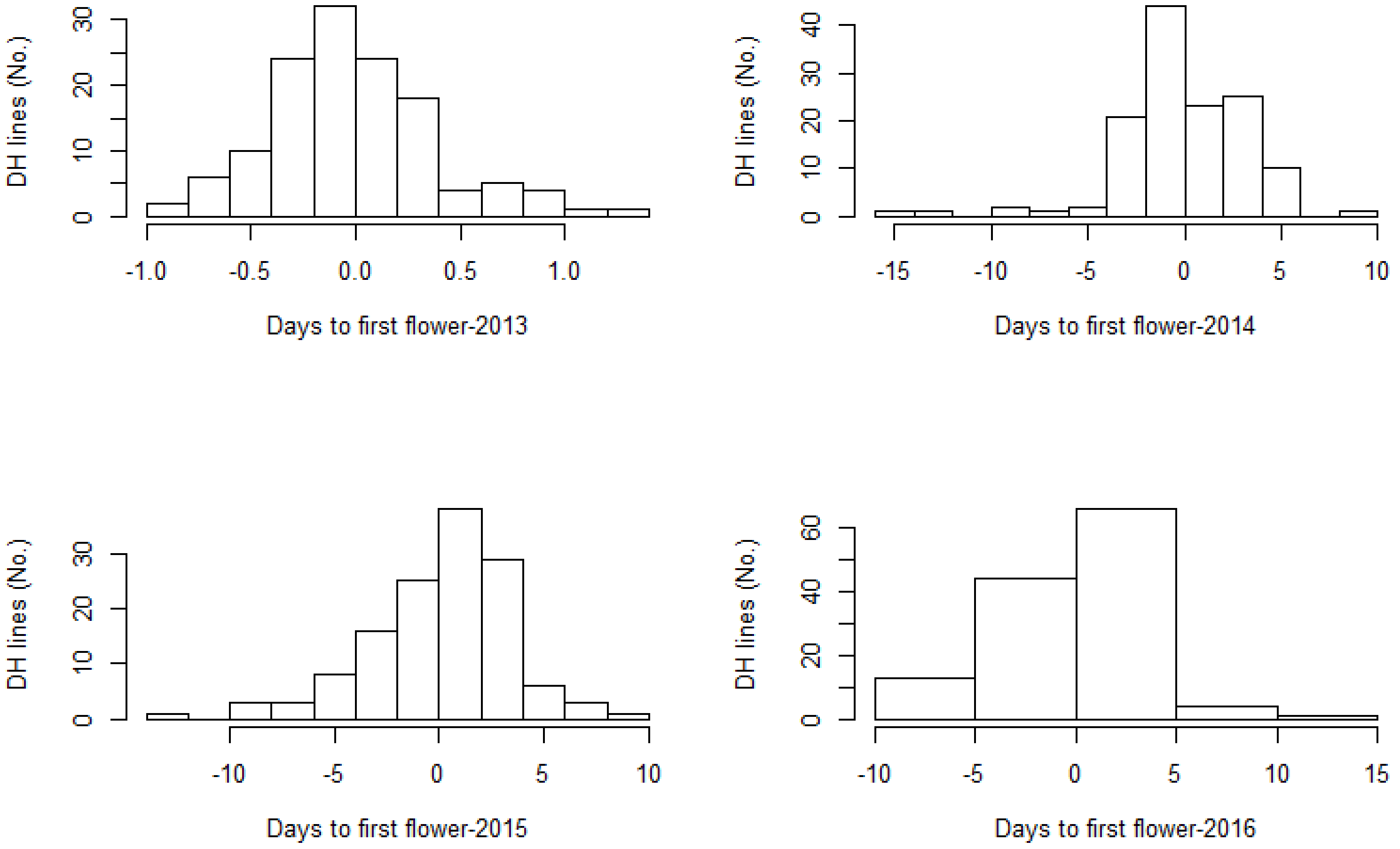

Supplemental Figure 6 (D). Frequency distribution of plant height and plant emergence in a SAgS DH population phenotyped in 2016 growing environments.

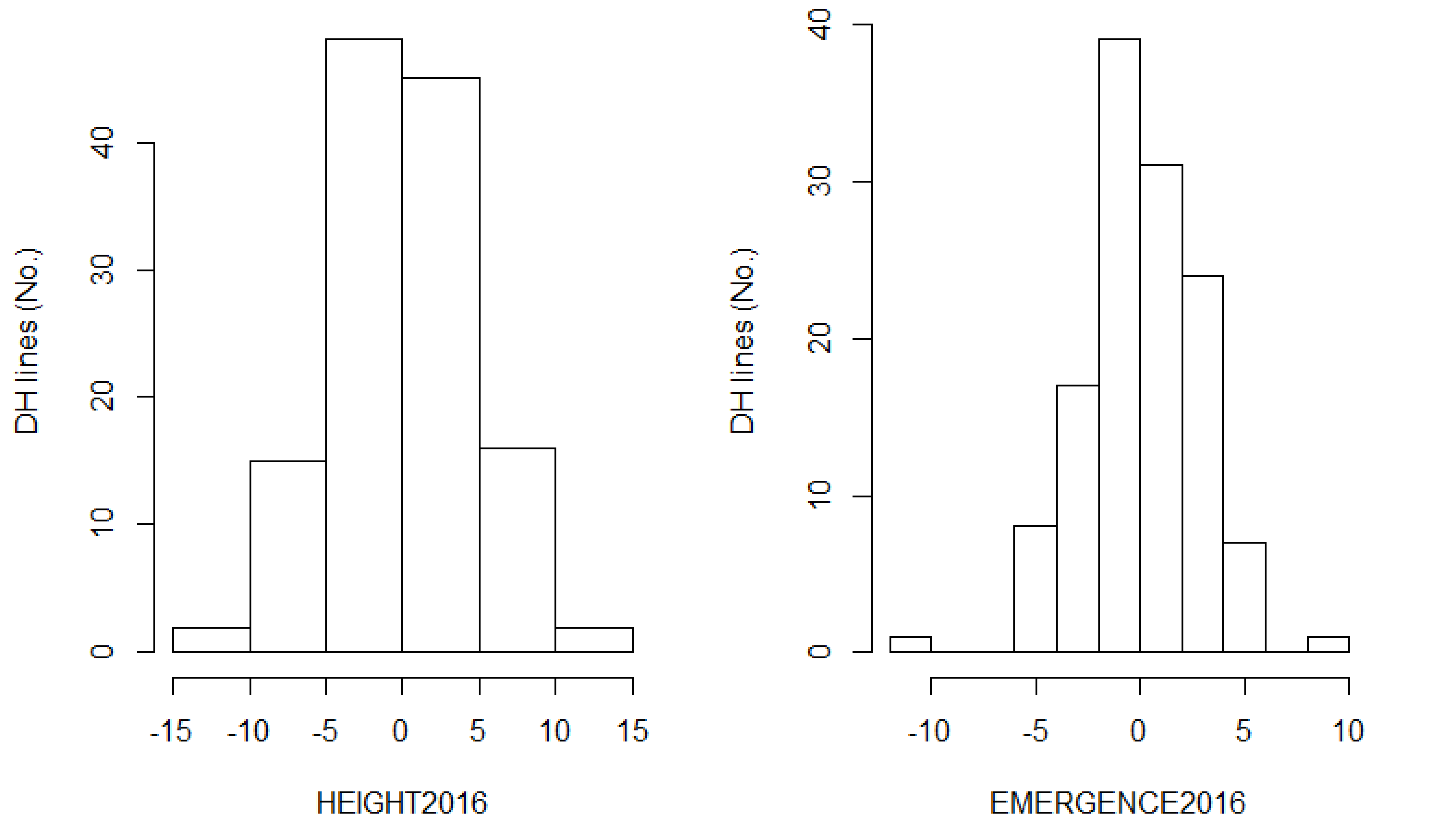

Supplemental Figure 6 (E). Frequency distribution of grain yield in a SAgS DH population phenotyped across four environments (2013-2016). Phenotypic data of 2013 and 2014 experiments was published previously (Raman et al 2016).

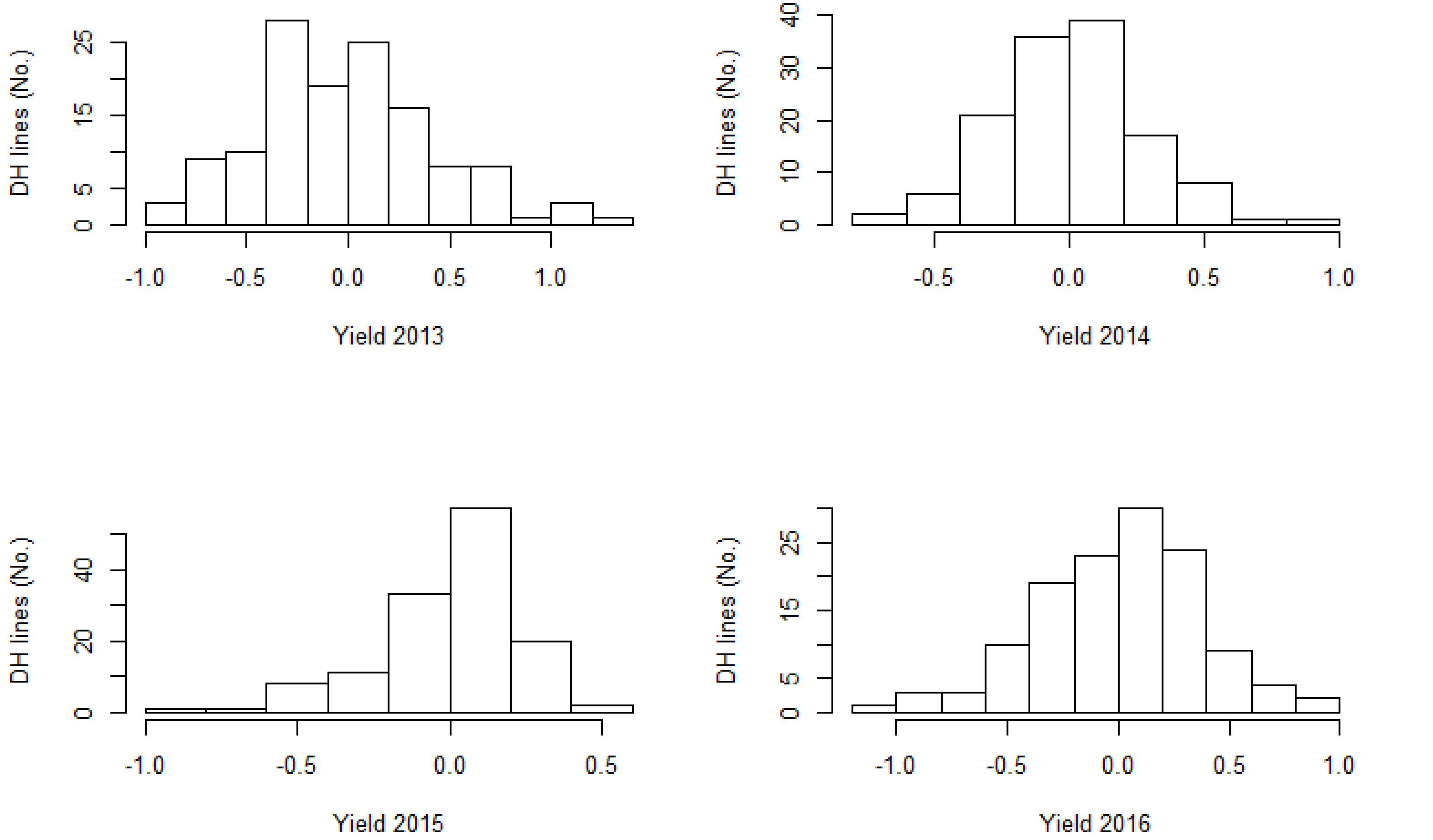
